# A nucleation barrier spring-loads the CBM signalosome for binary activation

**DOI:** 10.1101/2022.01.28.477912

**Authors:** Alejandro Rodríguez Gama, Tayla Miller, Jeffrey J. Lange, Jay Unruh, Randal Halfmann

**Affiliations:** Stowers Institute for Medical Research, Kansas City, Missouri; Department of Molecular and Integrative Physiology, University of Kansas Medical Center, Kansas City, Kansas

## Abstract

Immune cells activate in a binary, switch-like fashion that involves proteins polymerizing into large complexes known as signalosomes. The switch-like nature of signalosome formation has been proposed to result from large energy barriers to polymer nucleation. Whether such nucleation barriers indeed drive binary immune responses has not yet been shown. Here, we employed an in-cell biophysical approach to dissect the assembly mechanism of the CARD-BCL10-MALT1 (CBM) signalosome, a key determinant of transcription factor NF-κB activation in both innate and adaptive immunity. We found that the adaptor protein BCL10 encodes an intrinsic nucleation barrier, and that this barrier has been conserved from cnidaria to humans. Using optogenetic tools and a single-cell transcriptional reporter of NF-κB activity, we further revealed that endogenous human BCL10 is supersaturated even in unstimulated cells, indicating that the nucleation barrier operationally stores energy for subsequent activation. We found that upon stimulation, BCL10 nucleation by CARD9 multimers triggers self-templated polymerization that saturates NF-κB activation to produce a binary response. Pathogenic mutants of CARD9 that cause human immunodeficiencies eliminated nucleating activity. Conversely, a hyperactive cancer-causing mutation in BCL10 increased its spontaneous nucleation. Our results indicate that unassembled CBM signalosome components function analogously to a spring-loaded mousetrap, constitutively poised to activate NF-κB through irrevocable polymerization. This finding may inform our understanding of the root causes and progressive nature of pathogenic and age-associated inflammation.

## Introduction

Cells of both the innate and adaptive immune systems respond immediately and decisively to danger through the formation of large cytosolic protein complexes, known as signalosomes. Signalosomes function to coordinate the detection of pathogen- or danger-associated molecular patterns with protective transitions in cell state, such as programmed cell lysis or activation of nuclear transcription factor-κB (NF-κB) (Kellogg et al., 2015; Liu et al., 2014; Matyszewski et al., 2018; Wu and Fuxreiter, 2016). NF-κB directs the transcription of genes encoding pro-inflammatory cytokines and growth factors that are essential to both innate and adaptive immune responses, and its improper regulation contributes to cancer, chronic inflammation, and autoimmune diseases. Although NF-κB s known to respond in switch-like fashion to certain stimuli (Kingeter et al., 2010; Muñoz et al., 2019; Tay et al., 2010) (Fig 1A), the underlying positive feedback mechanism(s) responsible for this phenomenon remains unclear.

**Figure 1.**
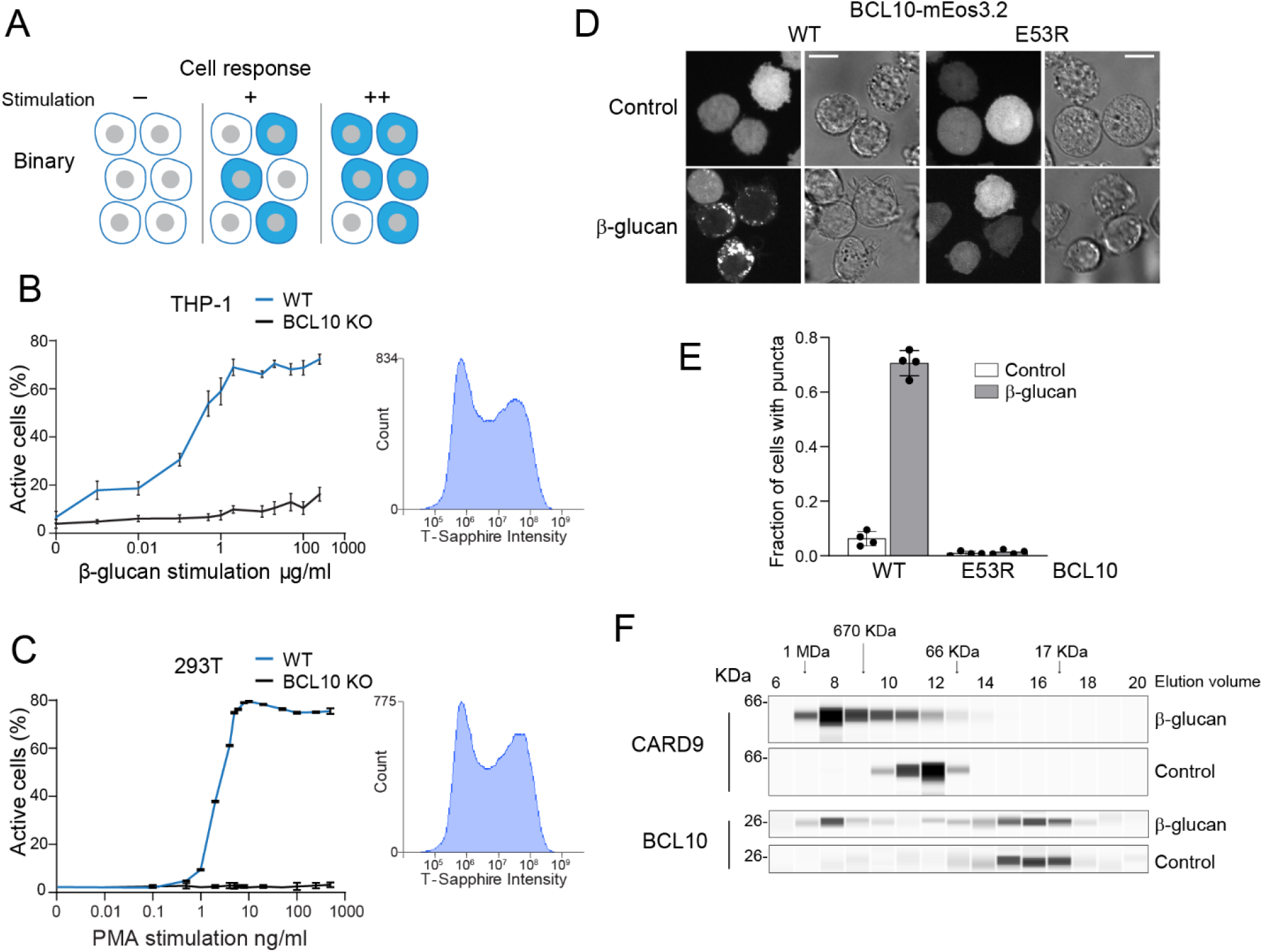
Assembly of the CBM signalosome drives all-or-none activation of NF-κB. (A) Schematic of binary activation in a population of cells. Higher doses of stimulation increase the probability of all-or-none NF-κB activation. (B) β-glucan stimulation of THP-1 WT and *BCL10*-KO monocytes transduced with an NF-κB transcriptional reporter. Monocytes were stimulated for 24 hrs then measured for T-Sapphire expression via flow cytometry. The graph shows the fraction of cells positive for NF-κB expression. Inset shows the histogram of T-sapphire expression indicating the distinct negative and positive population of THP-1 WT cells at stimulation of 10 μg/ml of β-glucan. (C) HEK293 WT and *BCL10*-KO cells were transduced with an NF-κB transcriptional reporter and stimulated with PMA for 24 hrs then measured for T-Sapphire expression via flow cytometry. Inset shows the expression of T-sapphire in HEK293T WT cells treated with 5 ng/ml of PMA. (D) THP-1 *BCL10*-KO monocytes were transduced with a doxycycline-inducible *BCL10*-mEos3.2 construct. Cells were imaged for BCL10 expression 24 hrs after Dox induction. Images show BCL10 puncta only in cells treated with β-glucan for 16 hrs (see also figure S2). In contrast, BCL10 E53R does not form puncta regardless of stimulation. Scale bar 10 μm. (E) Quantification of the number of THP-1 cells with BCL10 puncta after β-glucan stimulation. Each value represents the average of 4 independent experiments with more than 50 cells each. (F) Endogenous untagged CARD9 and BCL10 form high molecular weight species in THP-1 cells stimulated with β-glucan for 24 hrs. After treatment, cells were lysed and protein extracts were resolved by size exclusion chromatography followed by capillary immunodetection.

The CBM signalosome is a major determinant of NF-κB activation in lymphoid, myeloid, and certain non-immune cell lineages. Named for its three protein constituents, the CBM signalosome is composed of: 1) a Caspase Activation and Recruitment Domain (CARD)-coiled coil (CC) family member (either CARD9, 10, 11, or 14); 2) B cell lymphoma 10 (BCL10); and 3) mucosa-associated lymphoid tissue lymphoma translocation protein 1 (MALT1) (Gehring et al., 2018; Ruland and Hartjes, 2019; Ruland et al., 2001). The CARD-CC proteins have tissue-specific expression that places CBM formation under the control of specific cell surface receptors. CARD11 expression in the lymphoid lineage controls B and T cell activation upon antigen recognition by B- and T-cell receptors (Wang et al., 2002) CARD9 expression in the myeloid lineage links the antifungal response of monocytes to fungal carbohydrate detection by C-type lectin receptors (Gross et al., 2006; Hsu et al., 2007; Strasser et al., 2012). CARD10 and CARD14 are expressed in nonhematopoietic cells including intestinal and skin epithelia, respectively. Multiple clinically significant mutations in CARD proteins have been found to compromise immune system homeostasis. For instance, mutations that relieve the normally autoinhibited state of CARD11 promote lymphoid cell proliferation leading to lymphoma (Compagno et al., 2009; Lenz et al., 2008). Several mutations in CARD9 cause familial hypersusceptibility to fungal infections (Glocker et al., 2009; Lanternier et al., 2015, 2013). CARD10 mutations are associated with risk of primary open angle glaucoma (Zhou et al., 2016), while CARD14 gain of function mutations cause psoriasis (Howes et al., 2016; Jordan et al., 2012). Once activated, CARD-CC proteins recruit the adaptor protein BCL10 and its binding partner, MALT1, resulting in the activation of downstream effectors that ultimately allow NF-κB to translocate to the nucleus and induce transcription of its target genes. As the integrators of cell type-specific CARD-CC signaling, BCL10 and MALT1 are expressed ubiquitously across cell types. Point mutations and translocations involving BCL10 and MALT1 cause immunodeficiencies (Ruland and Hartjes, 2019), testicular cancer (Kuper-Hommel et al., 2013), and lymphomas (Zhang et al., 1999).

Bistability in proinflammatory pathways could result from positive feedback inherent to the process of signalosome assembly (Rodríguez Gama et al., 2021). This hypothesis draws from discoveries that certain signalosome adaptor proteins, including BCL10, are capable of self-templated or “prion-like” assembly in vitro or when overexpressed (Cai et al., 2014; Franklin et al., 2014; Holliday et al., 2019; Hou et al., 2011; Latty et al., 2018; Lu et al., 2014; Mompeán et al., 2018; O’Carroll et al., 2020; Qiao et al., 2013). Prion-like behavior is made possible by a structurally-encoded kinetic barrier to de novo assembly, a process known as “nucleation” (Khan et al., 2018; Rodríguez Gama et al., 2021). Nucleation involves a rare spontaneous acquisition of order from disorder. The nucleation barrier describes the improbability of that fluctuation occurring in a given window of time and space, and allows protein concentrations to locally exceed their thermodynamic solubility limits, i.e. become supersaturated. For prion-like signalosome proteins whose activity is coupled to assembly, sufficiently large nucleation barriers could allow resting cells to accumulate a driving force for subsequent abrupt activation, in response to a nucleating template that forms upon exposure to pathogens or cellular damage. For this mechanism to drive binary cellular decisions would require a) that the proteins retain the nucleation barrier in vivo; b) that they are supersaturated even at endogenous intracellular concentrations; and c) that polymerization can be triggered solely by the introduction of a template, i.e. is not driven by a reduction in thermodynamic solubility brought about by post-translational modifications or binding partners. These criteria have not yet been assessed for any signalosome.

In this study, we used a single cell reporter of NF-κB activity to reveal that the CBM signalosome activates NF-κB in a binary fashion. We then used our recently developed biophysical approach, Distributed Amphifluoric FRET (DAmFRET; (Khan et al., 2018; Venkatesan et al., 2019)), to dissect the mechanism of CBM signalosome formation. This effort uncovered a structurally-encoded nucleation barrier specifically in the adaptor protein, BCL10. We found that BCL10 polymer templates form within pre-existing CARD-CC protein multimers through a stimulus-dependent reorientation of their CARD domains. We further developed an optogenetic approach to nucleate BCL10 independently of upstream factors, revealing that BCL10 is supersaturated and poised to drive its own polymerization even in resting cells. Finally, we showed that the nucleation barrier of BCL10 is conserved from cnidaria to humans. Altogether, our work reveals that BCL10 functions as a nucleation-limited switch that makes cell decisions.

## Results

### Assembly of the CBM signalosome drives all-or-none activation of NF-κB

To explore the link between signalosome nucleation and signaling kinetics, we first developed a single-cell reporter of NF-κB activity (Fig S1A). This transcriptional reporter contains four copies of the core NF-κB response element followed by the coding sequence of the fluorescent protein, T-Sapphire (Wilson et al., 2013). We used human embryonic kidney 293T cells and THP-1 monocytic cells to measure the activation of NF-κB with increasing concentrations of CBM signalosome stimulation. To activate the CBM signalosome in THP-1 monocytes, we stimulated CARD9 using the yeast cell wall component, β-glucan. To activate the CBM signalosome in HEK293T cells, we stimulated CARD10 (the only CARD-CC expressed in HEK293T cells) using the PKC activator, PMA (Ruland et al., 2001). Using flow cytometry, we found that the percentage of T-Sapphire-positive cells increased in a dose-dependent manner for both HEK293T and THP-1 cells (Fig 1B and C). The distribution of T-Sapphire fluorescence was bimodal, with cells distributing between non-fluorescent and uniformly fluorescent populations. Increasing doses increased the fraction of cells in the fluorescent population, but did not influence the level of fluorescence within that population, even across multiple stimulation concentrations spanning several orders of magnitude (Fig 1B and C insets; Fig S1B and C). To ensure that activation was occurring solely through the CBM signalosome, we used CRISPR-Cas9 to disrupt exon 1 of *BCL10* in both HEK293T and THP-1 cells (Fig S2A-C). As expected, cells lacking BCL10 failed to respond to any stimulation (Fig 1B-C).

This all-or-none transition from inactive to active NF-κB at the cellular level implies the existence of positive feedback in the activity of one or more components of the signaling pathway (Fig 1A). To determine if positive feedback occurred at the level of NF-κB, we used TNF-α to activate NF-κB independently of the CBM signalosome. In contrast to our results with β-glucan and PMA, TNF-α activated NF-κB in a graded fashion. That is, T-Sapphire levels in individual cells increased with dose (Fig S1D and E). This result indicates that positive feedback occurs upstream of NF-κB.

We next investigated CBM signalosome assembly in THP-1 monocytes. To do so, we performed size exclusion chromatography of lysates from THP-1 monocytic cell lines either with or without β-glucan stimulation, and then immunoblotted for both BCL10 and CARD9. We found that stimulation caused CARD9 (62 kD) to shift uniformly to a higher molecular weight (Fig 1F). Bcl10 (26 kD) also populated a large (>500 kD) complex upon stimulation, although the size distribution was bimodal and a fraction of the protein remained monomeric. This suggests that Bcl10 but not CARD9 assembly occurs in a highly cooperative fashion that could, in principle (Koch, 2020), underlie the feed forward mechanism.

To determine if the high molecular weight assemblies require BCL10 polymerization, we knocked out *BCL10* in THP-1 cells and reconstituted it with either WT or polymerization-deficient mutant (E53R; (David et al., 2018)) BCL10-mEos3.2 expressed from a doxycycline-inducible promoter (Fig S2D). We then imaged cells following doxycycline addition, either with or without β-glucan treatment. In untreated cells, BCL10 fluorescence was entirely dispersed, consistent with the expected soluble form of the protein (Fig 1D). Conversely, in cells treated with β-glucan, WT BCL10 formed puncta in 65% of cells (Fig 1E), while the mutant protein remained dispersed. Together, these results suggest that the CBM signalosome, and particularly BCL10, via polymerization, exerts binary control over NF-κB activation in monocytes.

### The adaptor protein BCL10 is a nucleation-mediated switch

We next sought to understand the mechanism of positive feedback in CBM signalosome formation. Positive feedback in certain other innate immune signalosomes has been attributed to prion-like self-templated polymerization of death fold proteins (Cai et al., 2014; Franklin et al., 2014). CBM has a death fold domain in each of the three proteins -- a CARD domain in CARD9/10/11/14, a CARD domain in BCL10, and a death domain in MALT1 (Fig 2A). To determine whether one or more of these polymerizes in a switch-like fashion, we used DAmFRET to detect nucleation barriers for the full length proteins (Khan et al., 2018; Venkatesan et al., 2019). In this method, a protein of interest is tagged with the photoconvertible fluorescent protein mEos3.1 and expressed over a range of concentrations in cells. Upon photoconversion, molecules that are photoconverted act as FRET acceptors to the fluorophore molecules that have not converted. Protein self-assembly increases FRET signal, which is evaluated as a function of the protein’s concentration in each cell. We chose yeast cells as the reaction vessel because they contain neither death folds nor the associated signaling networks, which could otherwise obscure the exogenous proteins’ sequence-encoded assembly properties. We found that neither CARD9 nor MALT1 formed self-templating polymers. The former formed low AmFRET multimers across the full range of concentrations (Fig 2B). These corresponded to the protein’s accumulation into a single irregular punctum in each cell (Fig S3A). There was no discontinuity in DAmFRET and hence no apparent nucleation barrier to the formation of these puncta. MALT1 remained completely monomeric (Fig 3D, and Fig S3C). BCL10, in contrast, produced a clear bimodal distribution wherein cells partitioned between a no-AmFRET population and a high-AmFRET population, respectively comprised of cells containing only monomeric BCL10 or highly polymerized BCL10 (Fig 2C and Fig S3B). The two populations overlapped in their expression levels, indicating that BCL10 can accumulate to supersaturating concentrations while remaining monomeric, and then polymerizes following a stochastic nucleation event. The paucity of cells between the two populations, i.e. with moderate levels of AmFRET, suggests that BCL10 polymerizes rapidly to a new steady state following the nucleation event (Khan et al., 2018; Posey et al., 2021).

**Figure 2.**
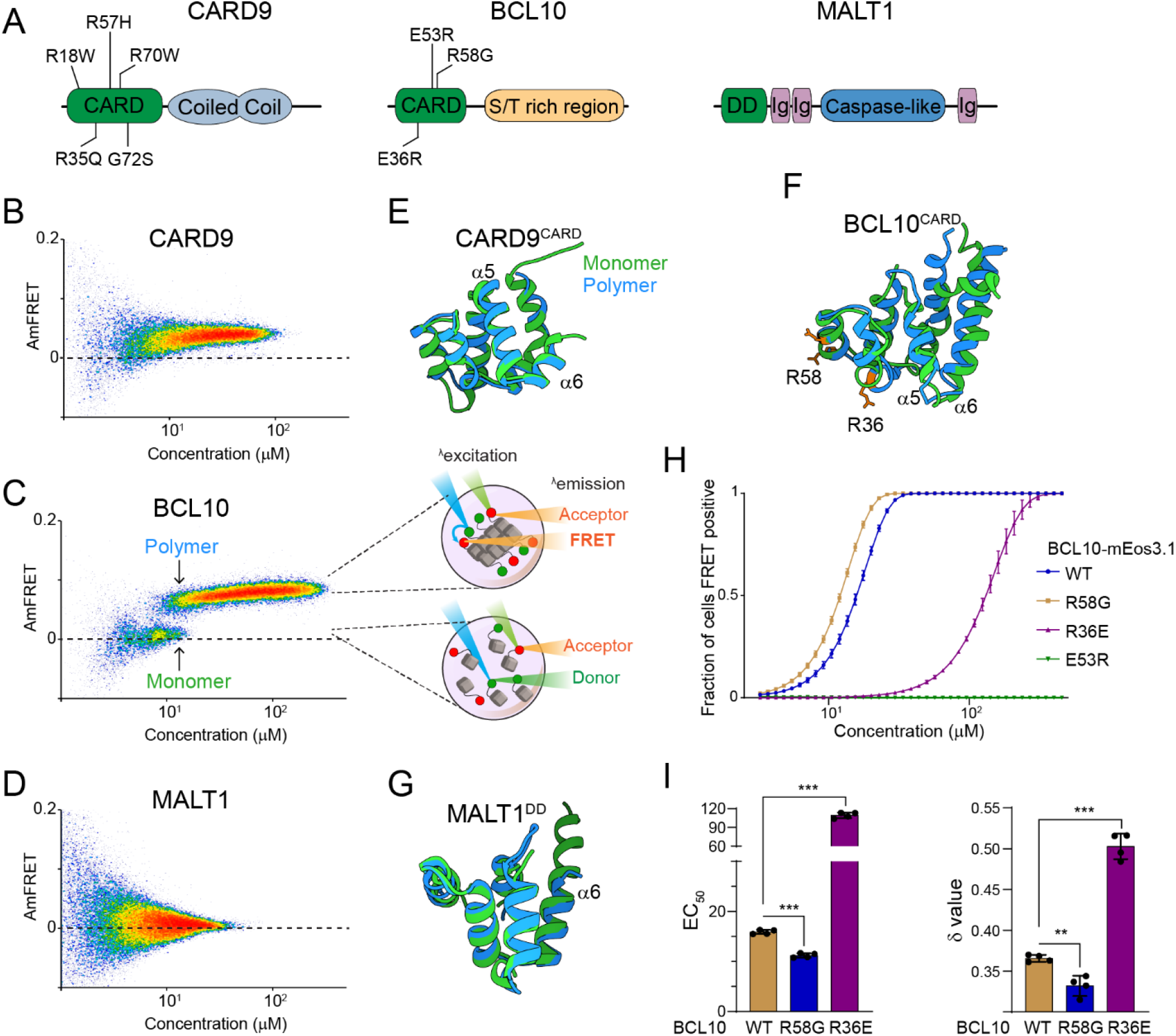
The adaptor protein BCL10 is a nucleation-mediated switch. (A) Domain architecture of CBM component proteins. Death fold domains are colored green. Mutations discussed in the text are indicated. (B) DAmFRET plot of CARD9 expressed in yeast cells, showing moderate AmFRET and therefore oligomer formation at all concentrations. (C) Left, DAmFRET plot of BCL10 expressed in yeast cells, showing a bimodal distribution between cells containing only monomers and cells containing polymers. The discontinuity and overlapping range of protein concentration for the two populations indicates that polymerization is rate-limited by a nucleation barrier. Right, schema of cells in the two populations, showing how self-assembly of partially photoconverted mEos3.1-fused proteins causes a FRET signal. (D) DAmFRET plot of MALT1 expressed in yeast cells, showing no AmFRET and therefore a monomeric state of the protein at all concentrations. (E) Structural alignment of CARD9^CARD^ monomers in its soluble (6E26, green) or polymerized (6N2P, blue) forms, showing no major conformational differences between them. (F) Structural alignment of BCL10^CARD^ monomers in its soluble (2MB9, green) or polymerized (6BZE, blue) forms, showing multiple conformational differences that span a-helices 1, 2, 5 and 6. (G) Structural alignment of MALT1^DD^ monomers in its soluble (2G7R, green) and polymerized (together with BCL10, 6GK2, blue) forms, showing no major conformational differences. (H) Weibull fits to DAmFRET plots of WT and mutant BCL10, showing how each mutation affects polymerization as quantified in (I). (I) EC_50_ (left) and □ (right) values derived from the fits in (H). EC_50_ describes the median concentration at which nucleation occurs. The slope parameter □ is a dimensionless proxy for the conformational free energy of nucleation. The R36E and E53R mutations, which lie in separate polymer interfaces, prevented the protein from polymerizing at low concentrations. The cancer-associated R58G mutation decreased the nucleation barrier, resulting in more frequent spontaneous nucleation relative to the WT protein. *** P<0.001, ** P<0.005

**Figure 3.**
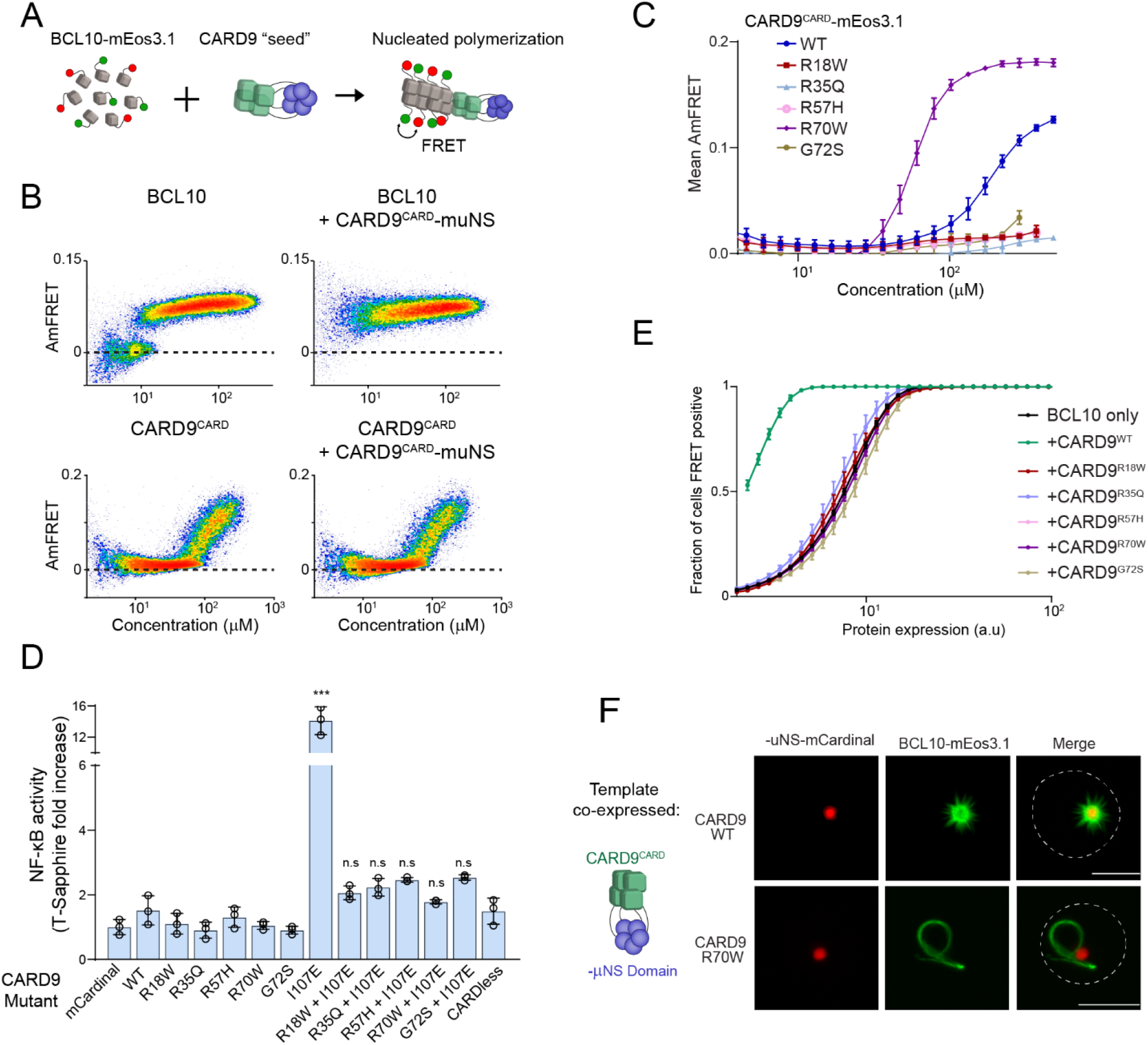
Insight into the mechanism of nucleation by CARD proteins and its disruption by pathogenic mutations. (A) Schematic of the DAmFRET experiment used to test for nucleating interactions between CBM components. Artificial “seeds” are created by expressing the putative nucleating protein as a fusion to the homo-oligomerizing module, μNS (purple). (B) DAmFRET plots of BCL10 (top) or CARD^CARD^ (bottom) in the absence (left) or presence (right) of CARD9^CARD^ seeds expressed in trans. For BCL10, the presence of seeds shifted all low-AmFRET cells to high-AmFRET, indicating that the protein had been supersaturated and is nucleated by CARD9^CARD^. In contrast, the seeds had no effect on CARD9^CARD^ DAmFRET, confirming that CARD9^CARD^ only polymerizes at high concentrations and does not become supersaturated. (C) Graph of spline fits of AmFRET values for CARD9^CARD^ mutants. All of the pathogenic mutations within the CARD domain disrupted polymer formation except for R70W, which stabilized polymers. (D) Effect of CARD9 mutations in the full-length protein context and with auto-repression eliminated by the I107E mutation. 293T transduced with the NF-κB reporter were transfected with the indicated CARD9-mCardinal constructs and analyzed for T-Sapphire expression at 48 hrs post-transfection. T-Sapphire expression was normalized to that of cells transfected with plasmid expressing mCardinal alone. ***P<0.0001 (E) Weibull fits to DAmFRET plots of BCL10 co-expressed with mutant CARD9 seeds, showing that only WT CARD9 nucleates BCL10 while none of the pathogenic mutants do so. (F) Images of yeast cells expressing the respective CARD9 seeds (red punctum) and BCL10-mEos3.1 (green). CARD9 WT seeds nucleate BCL10 polymers, resulting in a starburst-like structure. Conversely, CARD9 mutant R70W does not nucleate BCL10. BCL10 instead polymerizes spontaneously in these cells, resulting in a single long filament. Scale bar 5 μm.

We next evaluated nucleation barriers of the isolated death fold domains of CARD9, BCL10 and MALT1. MALT1^DD^ remained monomeric at all concentrations (Fig S3F). CARD9^CARD^ was monomeric up to a threshold concentration of approximately 100 uM, above which it readily polymerized with only a slight discontinuity (Fig S3D). In contrast, BCL10^CARD^ switched from monomer to polymer with a large discontinuity (Fig S3E) resembling that of full-length BCL10, suggesting that the large nucleation barrier of the full-length protein, and by extension the positive feedback associated with CBM assembly, can be attributed to the CARD domain of BCL10.

The structural basis of nucleation barriers that allow for certain death fold domains to supersaturate in cells is not completely understood. We reasoned that, as for amyloid-based prions, it may result from a conformational change required for subunit polymerization. To investigate this possibility, we took advantage of previously solved structures for all three of the CBM death fold domains. By computationally superposing the structures of the monomer and polymer forms for each domain, and calculating the root mean square deviation (RMSD) between backbone C-alpha atoms, we found that BCL10^CARD^ undergoes a greater conformational change to polymerize than does either CARD9^CARD^ or MALT1^DD^ (Fig 2E-G; RMSD 3.581, 1.655, and 0.956 Å, respectively), consistent with the nucleation barrier for BCL10^CARD^ resulting from a particularly unfavorable conformational fluctuation.

If the monomeric fold of BCL10 functions to restrict nucleation, then mutations that relax that fold can be expected to pathogenetically activate the CBM signalosome. The R58G mutation of BCL10 was first described in a germ cell line tumor and shown to hyperactivate NF-κB (Willis et al., 1999). We tested via DAmFRET if this can be attributed to a reduction in the sequence-encoded nucleation barrier. The EC_50_ and (delta) statistics obtained by fitting DAmFRET to a Weibull function serve as crude proxies for the inter- and intramolecular free energies of nucleation, respectively (Fig S3G). More specifically, they describe the median concentration at which nucleation occurs, and the independence of nucleation on concentration, respectively, where conformationally-limited nucleation has higher values (Khan et al., 2018). Indeed, relative to WT, R58G reduced both and EC_50_ (Fig 2H and I). In contrast, mutation R36E, which disrupts polymer interface IIa, increased both and EC_50_ (Fig 2I). Consistent with previous observations, mutant E53R, which disrupts polymer interface IIIb, blocked nucleation at all tested concentrations (Fig 2H). We next investigated the evolutionary conservation of R58. We found via multiple sequence alignment that position 58 is an arginine in mammals but a glutamine in lamprey, cartilaginous fish and amphibians (Fig S4A). We therefore evaluated the nucleation barrier when R58 is substituted with Gln or with a hydrophobic residue of comparable size (Leu). As expected, R58Q resembled WT, whereas R58L lowered the nucleation barrier (Fig S4B and C). These data suggest that R or Q residues at position 58 functionally oppose nucleation and thereby prevent precocious activation of BCL10 in the germ cell lineage. Altogether, we conclude that the native monomeric structure of BCL10 creates a physiological nucleation barrier that allows for switch-like self-assembly and all-or-none NF-κB activation.

### Pathogenic mutations disrupt BCL10 template formation within CARD-CC multimers

CARD-CC proteins form oligomers that promote BCL10 polymerization (David et al., 2018; Holliday et al., 2019). Somewhat paradoxically, however, CARD-CC proteins are believed to reside in an oligomeric state even in the absence of stimulation. The active configuration is prevented from forming within these oligomers due to an inhibitory interaction between the CARD domain and the adjacent coiled coil region (Holliday et al., 2019; Sommer et al., 2005). That interaction is released by post-translational modifications induced by pathway activation, leading to CBM assembly. The very low affinity of CARD9^CARD^ for itself (Fig 2B and (Holliday et al., 2019)) suggests that nucleation is unlikely to occur at physiological concentrations without assistance from coiled coil-mediated multimerization. This in turn necessitates that the coiled coil interactions support a stoichiometry higher than the dimer that we had previously observed for CARD9^1-142^ (Holliday et al., 2019), as a dimer has too few interfaces to template the four-start helical polymer of BCL10 (Qiao et al., 2013). Our observation that FL CARD9 forms a punctum at all expression levels, whereas the CARD domain itself forms either monomers (at low expression) or polymers (at high expression), confirms an essential role of non-CARD interactions in both autoinhibition and higher-order assembly.

To further dissect the relationship between multimerization, autoinhibition, and nucleation, we first used DAmFRET to assess CARD9 multimerization in the absence of its CARD domain. Remarkably, this truncated protein aggregated just as robustly as FL (Fig S5B and C), suggesting that coiled coil interactions are predominantly if not entirely responsible for its multimerization in the inactive state. Next, we progressively truncated the non-CARD region of the protein (Fig S5A). We found that even the shortest variant tested (1-142), retaining only 44 residues of additional sequence beyond the CARD domain, multimerized at all concentrations (Fig S5B and C). Unlike longer versions of the protein, however, this variant populated a lower AmFRET state and exclusively soluble oligomers, consistent with the reduced valency of the truncated coiled coil region (Fig S5D). Importantly, none of these constructs formed the higher AmFRET state or fibrillar puncta that would be indicative of polymerization. This tight relationship between multimerization and inhibition can be rationalized by the previously solved structure of the autoinhibited dimer formed by the short variant, wherein each CARD domain packs against the broad face of the immediately adjacent dimeric coiled coil, making extensive contacts with *both* helices (Holliday et al., 2019).

We had previously demonstrated that point mutations in the coiled coil-CARD interface release autoinhibition (Holliday et al., 2019). We therefore asked if the best characterized such mutation, I107E, has any effect on multimerization. Relative to WT, the mutant I107E achieved higher AmFRET at low expression and rendered the punctum less spherical (Fig S5E and F). Together with the prior demonstration of functional activity, the present findings confirm that multimerization itself is not inhibitory to nucleation and that once autoinhibition is released, CARD-CARD interactions stabilize the aggregated state. We therefore speculate that pre-multimerization driven by the multivalent coiled coil expedites the response to stimulation, perhaps by allowing stimulus-dependent post-translational modifications (Zhong et al., 2018) to occur cooperatively on multiple CARD9 subunits in close proximity. This would reduce the entropic cost of CARD9 activation relative to the subunits oligomerizing de novo.

To test if a high local concentration indeed suffices for CARD9^CARD^ subunits to organize into BCL10 polymer nuclei, we substituted the coiled coil region with an inert and well-characterized homomultimeric domain, μNS (Schmitz et al., 2009). This domain forms stable condensates that drive its fusion partner to very high local concentration (Fig 3A), and thereby eliminates intermolecular entropic contributions to nucleation barriers (Kandola et al., 2021). Expressing the CARD9^CARD^-μNS fusion should force to high-AmFRET any cells expressing either CARD9^CARD^- or BCL10-mEos3.1 above their respective concentration thresholds for polymerization. As hypothesized, CARD9^CARD^-μNS shifted all BCL10-mEos3.1 cells to the high-FRET population (Fig 3B), confirming that CARD9^CARD^ indeed forms a polymer seed when condensed. Moreover, the data confirm that in the absence of a template, BCL10 is supersaturated even at the lowest levels of expression (nanomolar). In striking contrast, CARD9^CARD^-μNS had no effect on the DAmFRET profile of CARD9^CARD^-mEos3.1, confirming that CARD9^CARD^ instead forms labile polymers with no appreciable nucleation barrier (Fig 3B).

We next asked if the other CARD-CC family members -- CARD10, 11 and 14 -- function in the same fashion. These proteins have the same domain architecture as CARD9: a CARD domain followed by coiled coil domains. We performed DAmFRET on their CARD domains and found that all polymerized comparably to that of CARD9 (Fig S6A), i.e. only at high concentrations (the most stable polymers were formed by CARD10, but even then only above approximately 40 uM), and with negligible nucleation barriers. We then asked if they too, when artificially condensed by fusion to μNS, suffice to nucleate BCL10. Indeed all exhibited this activity (Fig S6B). These findings suggest a shared mechanism of action by all four CARD-CC proteins.

Several mutations in CARD9^CARD^ cause susceptibility to fungal infections in humans (Fig 2A). We noted that these residues are conserved within the CARD-CC proteins (Fig S7A). To determine their mode(s) of action, we first asked if the mutants can form the polymer structure using DAmFRET and fluorescence microscopy. Whereas WT CARD9^CARD^ assembled into polymers at high concentration, the pathogenic mutants R18W, R35Q, R57H, and G72S failed to do so (Fig 3C and S7B,C), explaining why they cannot nucleate BCL10. Remarkably, however, the R70W mutant not only retained the ability to polymerize but did so at lower concentrations than WT (Fig 3C and Fig S7B,C), implying that its defect instead lies downstream of polymer formation.

To investigate downstream consequences of the CARD domain mutations, and in the context of the full-length protein, we next transiently transfected our NF-κB reporter cells with full-length versions of CARD9, all harboring the I107E mutation to eliminate autoinhibition. We first confirmed that CARD9 I107E (absent the pathogenic mutations) strongly activated NF-κB, and that a construct lacking the CARD domain did not (Fig 3D). We next introduced each of the pathogenic mutants into this construct and found that every one of them eliminated the protein’s ability to activate NF-κB (Fig 3D). All variants expressed to similar levels as WT (Fig S7D). Because the R70W mutant fails to template BCL10 despite robust polymerization, we conclude that this mutation specifically disrupts the nucleating interaction with BCL10 at interface IIb (Fig 3E, F and S7E). Altogether, these results confirm that CARD9 and its orthologs function to nucleate BCL10 by forming a polymeric structure stabilized by multivalent coiled-coil interactions, and that disrupting that activity compromises innate immunity.

### The BCL10 nucleation barrier causes binary activation of NF-κB

We next asked if switch-like BCL10 polymerization results in binary activation of NF-κB in human cells. To do so, we monitored NF-κB activation with respect to polymerization by CBM signalosome components. We transfected constructs of proteins fused to mEos3.2 into HEK293T cells containing the transcriptional reporter (T-Sapphire) of NF-κB activation (Fig 4A). Using DAmFRET, we confirmed that all of the proteins behaved the same way in HEK293T cells as they did in yeast (Fig 4B). Specifically, full length CARD9 formed irregular higher order assemblies at all concentrations (Fig S8A and C); CARD9^CARD^ polymerized only above a high threshold expression level; MALT1 was entirely monomeric (Fig S8A); and BCL10 became supersaturated with respect to a polymerized form (Fig 4B). Neither CARD9 nor MALT1 expression activated NF-κB (Fig S8A and B). BCL10 expression, in contrast, robustly activated NF-κB (Fig 4C). Importantly, this activation only occurred in cells that contained BCL10 polymers, and not monomers, irrespective of the expression level of BCL10 (Fig 4C, S8D,E). The frequency of activation, but not the level of activation within individual cells, was reduced or eliminated for the polymer-inhibiting mutants R36E and E53R, respectively (Fig S8D and E). Mutant R58G increased the fraction of cells with activated NF-κB (Fig S9C), as expected from its reduced nucleation barrier. Importantly, the intensity of NF-κB response at the cellular level did not differ between the mutants (Fig S8E and S9A-C), confirming a specific effect on nucleation and not downstream signaling. To determine if activation was specific to BCL10 polymers, we also performed the experiment with ASC, the death fold-containing adaptor protein of a different signalosome (the inflammasome). As expected (Cai et al., 2014), ASC formed prion-like polymers (Fig 4B), but did not activate the NF-κB reporter (Fig 4C). This experiment confirms that polymers of BCL10, specifically, and independently of upstream components or physiological stimuli, suffice to activate downstream components of the pathway, and that an intrinsic nucleation barrier to their formation causes activation to occur in a binary fashion.

**Figure 4.**
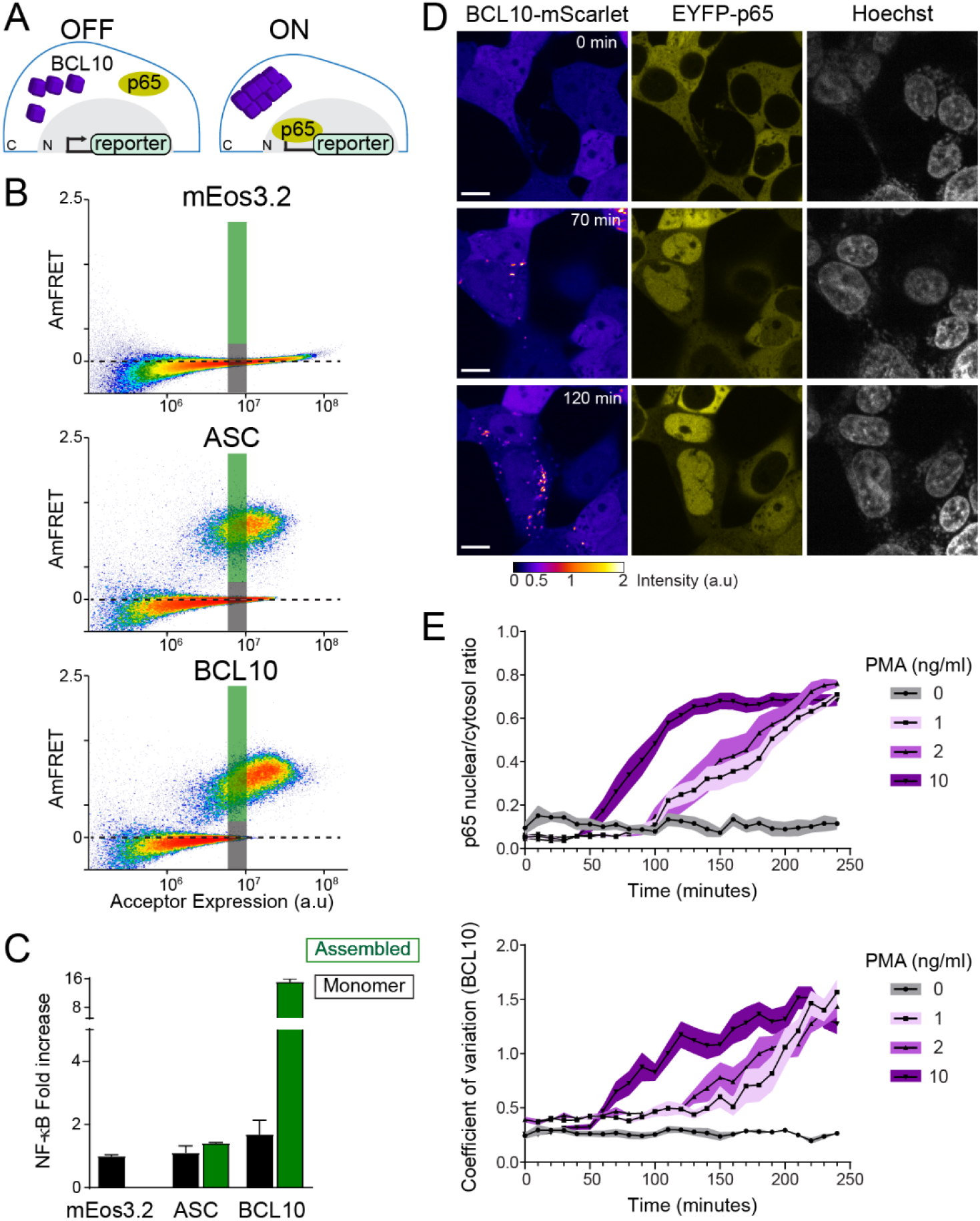
The BCL10 nucleation barrier causes binary activation of NF-κB. (A) Schematic of experiments to determine if BCL10 polymerization activates NF-κB in 293T cells. In the OFF state, BCL10 is dispersed in the cell and the p65 NF-κB subunit is localized to the cytosol. In the ON state, BCL10 is polymerized and p65 relocates to the nucleus, where it induces the transcription of NF-κB response genes. (B) DAmFRET plots of the indicated proteins expressed in 293T cells containing the NF-κB reporter, 48 hrs after transfection. The reporter mEos3.2 remains in the monomeric state and does not activate NF-κB. The gray and green boxes designate the regions gated for cells containing either monomer or polymerized protein, respectively, and analyzed for T-Sapphire expression in (C). (C) Quantification of T-Sapphire expression in the gated areas of the DAmFRET plots shown in (B). Only cells that contained BCL10 polymers, but not cells containing the same concentration of monomeric BCL10 nor cells containing ASC polymers, activated NF-kB. (D) Time-lapse microscopy of 293T *BCL10*-KO cells reconstituted with *BCL10*-mScarlet and EYFP-p65. Cells were stimulated with 10 ng/ml PMA and imaged every 10 min. BCL10 and p65 are distributed throughout the cytosol at 0 min, but respectively relocalize to cytosolic puncta and the nucleus over the course of the experiment. Scale bar 10 μm. (E) Top, ratios of EYFP fluorescence intensities in the nucleus versus cytosol; and bottom, values of the coefficient of variation of mScarlet pixel intensities, over the time course of cells stimulated with PMA at the indicated concentrations. BCL10 polymerization produced visible puncta within approximately 50 min of PMA stimulation. More than 40 cells were tracked and analyzed for each treatment condition.

### BCL10 is endogenously supersaturated

While the results obtained thus far indicate that BCL10 *can* function as a switch via nucleation-limited polymerization, we next asked whether it indeed does so even at endogenous levels of expression, i.e. in the absence of overexpression to promote BCL10 assembly. This distinction is very important because it determines if the nucleation barrier is responsible for preventing NF-κB activation in the absence of stimulation. To determine if the protein is *constitutively* supersaturated, we designed the experiment to eliminate any upregulation of *BCL10* transcription or translation that might occur upon stimulation (Yan et al., 2008). Specifically, we reconstituted our BCL10-deficient HEK293T cells with BCL10-mScarlet expressed from a doxycycline-inducible promoter. After isolating a single clone with uniform expression, we performed a doxycycline titration and used capillary protein immunodetection to compare the expression level of reconstituted BCL10-mScarlet to that of endogenous BCL10 in unstimulated HEK293T. We found that 1 ug/ml doxycycline induced BCL10-mScarlet to approximately the same level as endogenous BCL10 (Fig S10A-C). We next compared BCL10 expression levels in HEK293T cells to that of THP-1 cells and primary human fibroblasts, and found comparable expression levels across all three cell lines (Fig S10D and E). Finally, we analyzed BCL10 expression at the single cell level using immunofluorescence and flow cytometry, and confirmed that the reconstituted expression of BCL10-mScarlet has the same median intensity of BCL10 antibody binding as that of endogenous BCL10 in unmodified HEK293T cells (Fig S10F).

This cell line enables us to monitor NF-κB activity with respect to the assembly state of BCL10 at endogenous unstimulated levels of expression. To achieve this, we introduced a fluorescently tagged NF-κB subunit, EYFP-p65, and performed time lapse fluorescence microscopy following the addition of PMA. Prior to stimulation, both BCL10 and p65 were diffusely distributed throughout the cytosol. BCL10 became punctate within 50 minutes after PMA addition, and the puncta could be observed to elongate with time (Fig 4D, E and Video 1). Concomitantly, p65 was observed to translocate to the nucleus (Fig 4E). This result suggests that BCL10 is indeed supersaturated prior to stimulation.

To definitively test this possibility, we next devised an optogenetic approach to nucleate BCL10 independently of any possible reduction in its thermodynamic solubility that could come about through a cellular response to physiological stimuli, such as post-translational modifications or changes in the binding of other proteins to BCL10. Cry2clust is a variant of a plant photoreceptor that reversibly homo-oligomerizes in response to blue light excitation (Park et al., 2017). We expressed CARD9^CARD^ as a fusion to miRFP670-Cry2clust (CARD9^CARD^-Cry2) in the BCL10-mScarlet cell line (Fig 5A), and used a brief (1s) pulse of 488 nm laser light to activate Cry2. Prior to blue light exposure, both CARD9^CARD^-Cry2 and BCL10-mScarlet were diffusely distributed throughout the cells (Fig 5B and Video 2). Following the pulse, however, both proteins assembled into puncta. By measuring the kinetics of CARD9^CARD^ and Bcl10 assembly (as the coefficient of variation in pixel intensity over time), we found that CARD9^CARD^ clustering peaked at approximately five minutes post-excitation, and then disassembled to background levels by approximately 20 minutes (Fig 5C). Within six minutes following the pulse, BCL10 began to form filamentous puncta that initially co-localized with those of CARD9^CARD^ (Fig 5B). Remarkably, the puncta continued to elongate even after the CARD9^CARD^ clusters had dissolved (Fig 5B and inset), confirming that BCL10 is thermodynamically driven to a polymeric state even in the absence of CARD9 templates. To test the specificity of the CARD9-BCL10 interaction, we also performed the experiment with the R18W and R70W mutants of CARD9^CARD^-Cry2, whereupon BCL10 did not polymerize despite comparable kinetics of CARD9^CARD^-Cry2 clustering (Fig S11 A-D).

**Figure 5.**
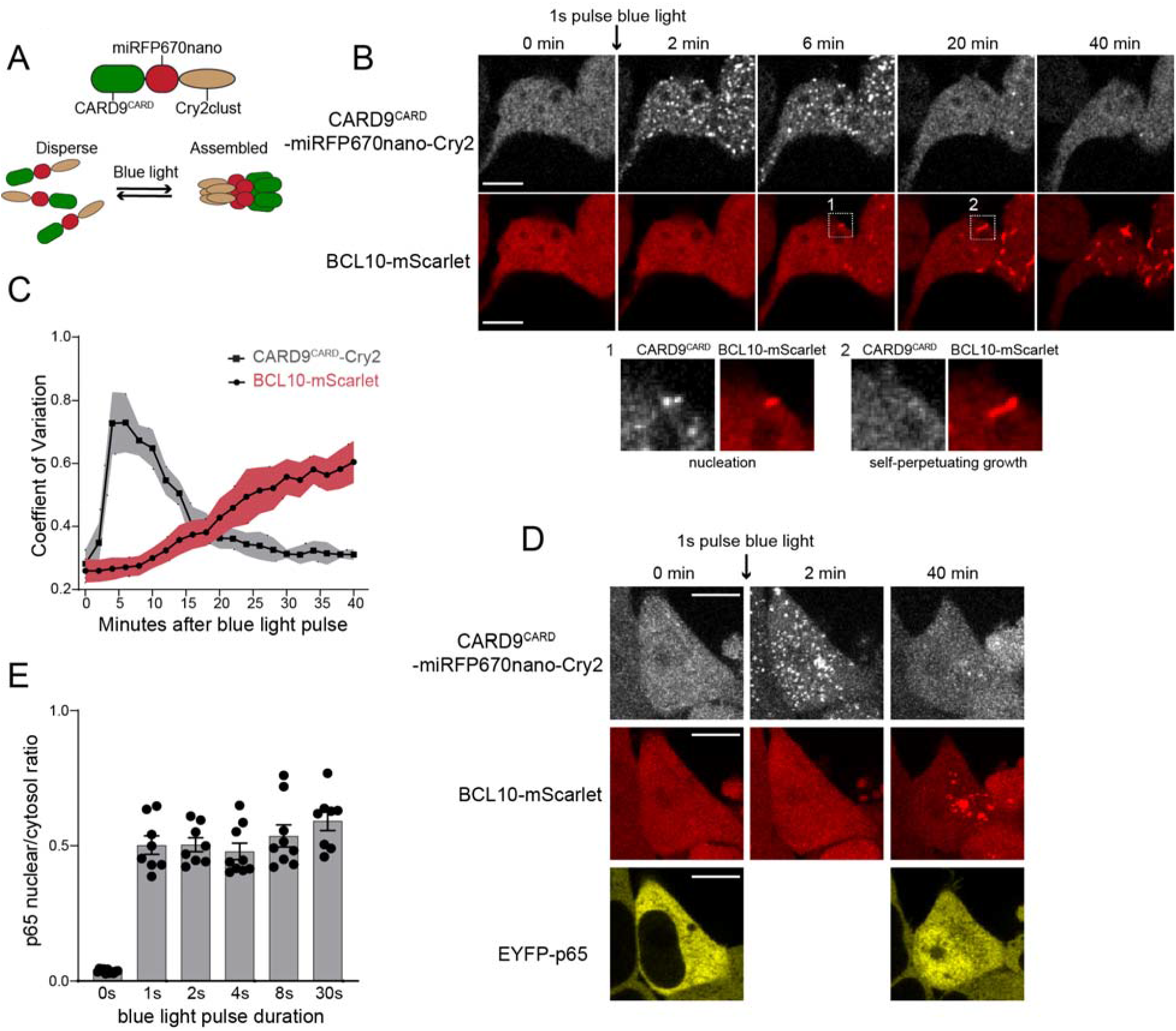
BCL10 is endogenously supersaturated. (A) Schematic of the optogenetic nucleation experiment. CARD9^CARD^ is expressed as a fusion to the near-infrared fluorescent protein miRFP670nano and the light-controlled homo-multimerizing module Cry2clust. Upon blue light exposure, Cry2clust forms reversible multimers that constrain the fused CARD9^CARD^ domains to a local concentration sufficient to form a polymeric template for BCL10. (B) Microscopy images of 293T *BCL10*-KO cells stably expressing CARD9^CARD^-miRFP670nano-Cry2clust and BCL10-mScarlet. Cells were maintained at 37°C with 5% CO2 and imaged every 2 minutes followed by a short pulse of 488nm light stimulation. Inset number 1, at 6 minutes post-stimulation, shows BCL10 colocalizing with a CARD9^CARD^ cluster. Inset number 2, at 20 minutes, shows that the CARD9^CARD^ cluster has dissolved, while the BCL10 punctum has instead elongated into a filament. Scale bar 10 μm. (C) Values of the coefficient of variation of mIRFP670nano and mScarlet pixel intensities over time. The data are from 40 cell measurements. (D) Microscopy images of cells expressing CARD9^CARD^ seeds, BCL10-mScarlet and EYFP-p65. Images were taken every 2 min following blue light stimulation. For technical reasons, EYFP-p65 was imaged only at the first and last time points. Scale bar 10 μm. (E) Ratios of EYFP fluorescence intensities in the nucleus versus cytosol after blue light stimulation for the indicated durations. Data are from at least 10 cells per treatment.

We then asked if the optogenetically-nucleated BCL10 polymers sufficed to induce the nuclear translocation of EYFP-p65. In order to avoid repeated stimulation of Cry2 oligomerization with the 514 nm laser, we only imaged EYFP-p65 at the beginning and end (40 min following the blue light pulse) of the experiment. We observed that blue light induced EYFP-p65 to translocate comparably to stimulation with PMA (Fig 5D and E). Additionally, as with the different doses of PMA, the degree of translocation within cells did not depend on the duration of the blue light pulse. This again highlights the all-or-none nature of signaling from BCL10. We confirmed that the polymerization-incompetent mutant of BCL10, E53R, failed to form puncta or activate NF-κB in response to blue light (Fig S12A-C). Altogether these data confirm that BCL10 is endogenously supersaturated and physiologically restrained from activation by the kinetic barrier associated with polymer nucleation, resulting in switch-like CBM signalosome formation and NF-κB activation.

### MALT1 activation depends on the multivalency rather than ordered structure of BCL10 polymers

That BCL10 function derives from a nucleation barrier implies that the structure of the active state has evolved toward increased order relative to the inactive state, irrespective of its interactions with downstream signaling components. If so, the well-ordered nature of BCL10 polymers (as distinct from merely a high density of subunits) should be irrelevant for MALT1 activation. To determine if there is any specific requirement of the BCL10 polymer structure for downstream signaling, we asked if MALT1 can be activated simply by increasing its local proximity in the absence of an ordered scaffold. To do so, we used optogenetic approaches to directly dimerize or multimerize MALT1 lacking its death fold domain. Specifically, we fused MALT1^126-824^ to either the blue light-dependent dimerization module, VfAU1-LOV, or the blue light-dependent multimerization module, Cry2clust, followed by the fluorescent protein miRFP670nano (Fig 6A and B). When expressed in HEK293T cells containing EYFP-p65 and illuminated with blue light, the dimerizing module failed to induce nuclear translocation, while the multimerizing module did so robustly, and to the same levels as had been achieved for BCL10-mediated activation (Fig 6A-D). As a control that the constructs were each functioning as intended, we also performed experiments with the analogous region of CASP8 (180-479) fused to either VfAU1-LOV or Cry2clust. CASP8 has previously been shown to activate upon dimerization (Demarco et al., 2020). Indeed, both constructs triggered rampant cell death upon blue light illumination (Fig S13A and B). Together these results reveal that MALT1 is activated by proximity rather than structural order of the adaptor protein polymers, consistent with a kinetic rather than thermodynamic function of BCL10 polymers. The fact that MALT1 activation requires higher order multimerization rather than just dimerization may relate to its functioning as a scaffold for the ubiquitin ligase TRAF6, a critical intermediary in the activation of NF-κB (Ginster et al., 2017).

**Figure 6.**
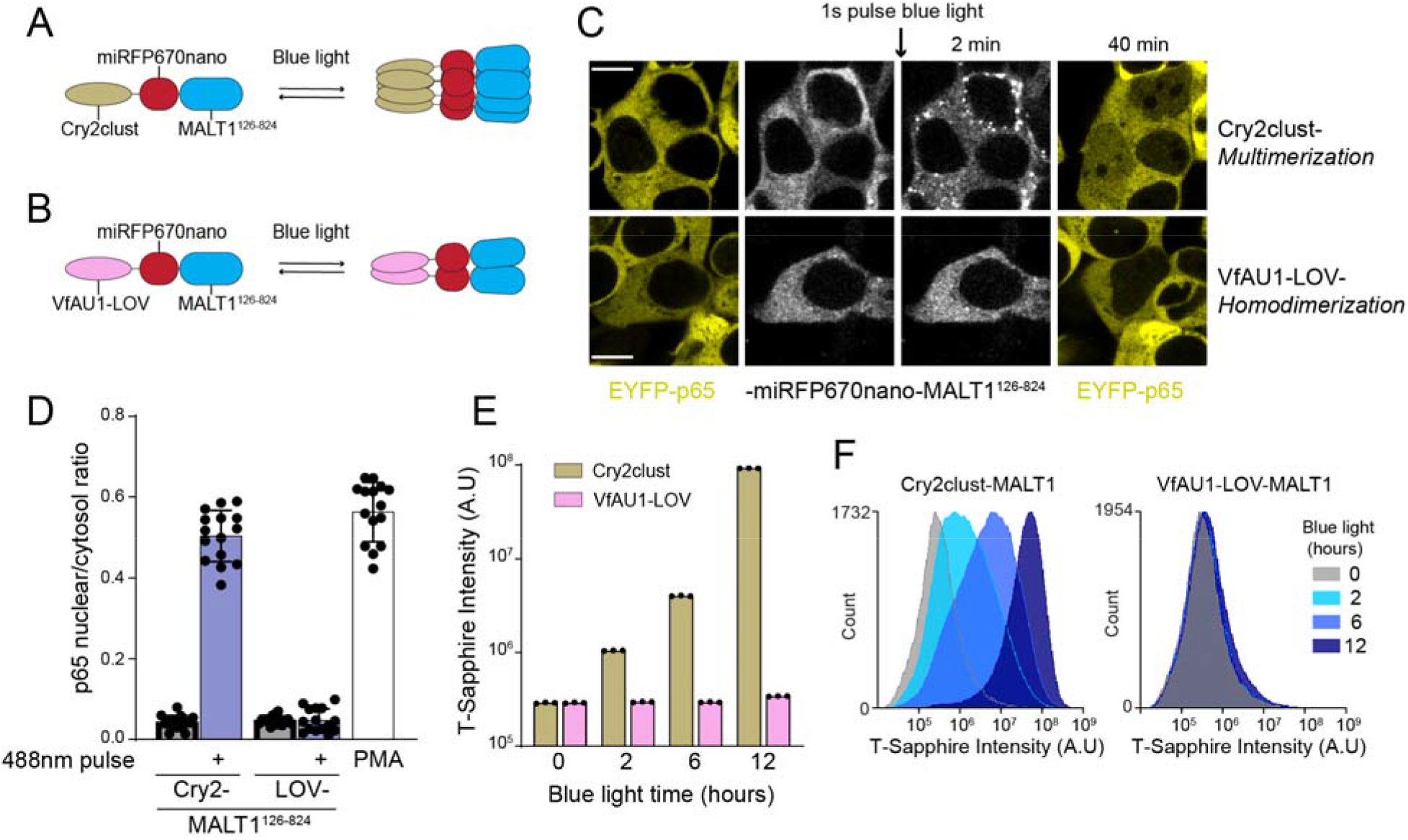
MALT1 activation depends on the multivalency rather than specific structure of BCL10 polymers. (A) Schematic of the optogenetic MALT1 activation experiment. A fragment of MALT1 lacking its death domain (MALT1^126-824^) is expressed as a fusion to mIRFP670nano and Cry2clust, allowing for its oligomerization upon blue light exposure. (B) Schematic of optogenetic MALT1 dimerization experiment. As for (A), but with Cry2clust replaced with the light-controlled dimerizing module, VfAU1-LOV. (C) Microscopy images of 293T cells expressing EYFP-p65 transfected with either the construct in (A) or (B). Images were taken every 2 min following a single pulse of blue light. EYFP-p65 was imaged before blue light stimulation and again 40 minutes after stimulation, showing that it translocated to the nucleus only in cells in which MALT1 had multimerized. Scale bar 10 μm. (D) Ratios of EYFP fluorescence intensities in the nucleus versus cytosol at the 40 minute time point for the experiments in (C). Data are from 15 cells per group. (E) NF-κB activation in 293T *MALT1-KO* NF-κB reporter cells transfected with the constructs in (A) and (B), and stimulated with blue light for the indicated durations. Data represent medians of three independent measurements. (F) Flow cytometry histograms of T-Sapphire fluorescence for the experiment in (E).

If the binary response of NF-κB to CBM stimuli is indeed conferred by BCL10, then it should switch to a graded response when MALT1 is activated optogenetically. To test this prediction, we expressed the Cry2clust- and VfAU1-LOV MALT1^126-824^ fusions in our NF-κB reporter cell line, and examined the population-level distribution of NF-κB activity in single cells exposed to blue light for different durations. As expected, the intensity of T-Sapphire fluorescence increased in a monotonic fashion with the duration of blue light stimulation (Fig 6E and F). This contrasts with the bimodal distribution that had been produced by physiological CBM stimulation and further illustrates that switch-like activation of NF-κB is conferred by BCL10.

### Ancient origin of the sequence-encoded nucleation barrier in BCL10

The CBM signalosome originated in the ancient ancestor to cnidarians and bilaterians (Staal et al., 2018) (Fig 7A). To determine if binarization is a conserved ancestral function of this signaling pathway, we investigated the structure and assembly properties of BCL10 from the model cnidarian, *Nematostella vectensis* (Nv). Using AlphaFold (Jumper et al., 2021), we found that the overall structure of the CARD domain, predicted with high confidence, is highly similar to that of human BCL10, and retains key residues of the polymer interface, such as E53 (Fig S14A-C).

**Figure 7.**
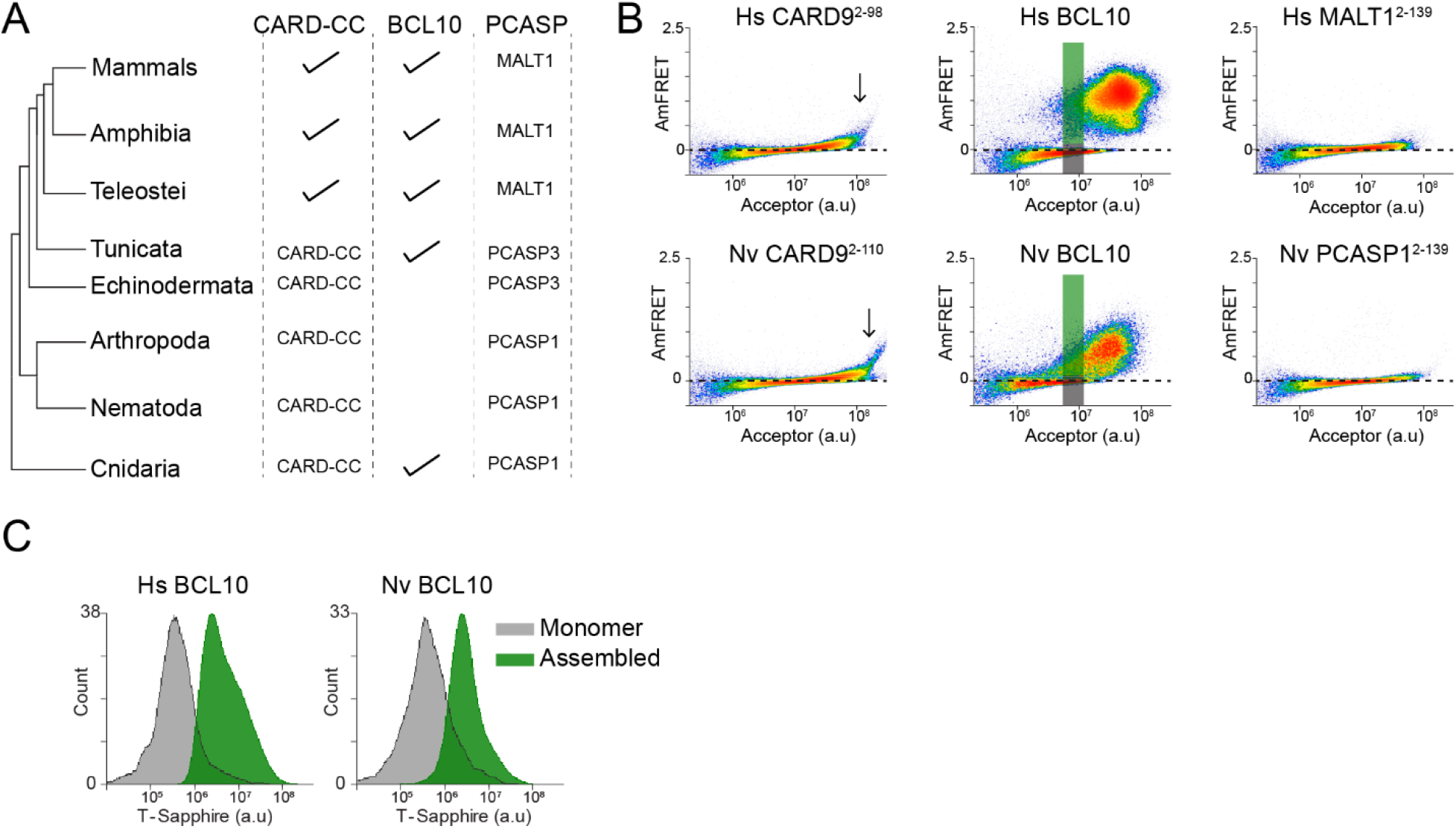
Ancient origin of the sequence-encoded nucleation barrier in BCL10. (A) Phylogenetic tree of CBM signalosome components in the major metazoan clades. One or more CARD-CC proteins occur in all clades, as does a MALT1-like paracaspase. BCL10 is found in chordates and cnidarians. (B) DAmFRET plots of 293T cells expressing CARD9^CARD^, BCL10, or MALT1^DD^ from human (top) or the sea anemone *Nematostella vectensis* (bottom). The gray and green boxes designate the regions gated for cells containing either monomer or polymerized protein, respectively, and analyzed for T-Sapphire expression in (C). (C) Flow cytometry histograms of T-Sapphire fluorescence in 293T *BCL10*-KO NF-κB reporter cells 48 hrs after transfection with either human or *N. vectensis BCL10*-mEos3.2. Gray and green shading represent cells containing either monomeric or polymerized BCL10, respectively, from the corresponding gates illustrated in (B). Data are representative of three independent experiments.

We next expressed each of the Nv CBM components as mEos3.2 fusions in human cells. We found that the DAmFRET profiles for all three proteins are strikingly similar to those of their human counterparts. Specifically, the CARD domain of CARD-CC, the CARD9 homolog, polymerized with a high saturating concentration and negligible nucleation barrier (Fig 7B). Nv BCL10 exhibited a discontinuous, nucleation-limited transition between monomer and polymer, and the death fold from Nv PCASP1, the MALT1 homolog, remained entirely monomeric (Fig 7B).

Finally, we asked if NvBCL10 can functionally replace human BCL10 with respect to binary activation of NF-κB. We expressed the protein in HEK293T *BCL10*-KO cells containing the transcriptional reporter (T-Sapphire) of NF-κB activation. By measuring T-Sapphire fluorescence within a DAmFRET experiment, we confirmed that indeed, the population of cells with NvBCL10 polymers also contained activated NF-κB, while cells expressing the same concentration of NvBCL10 in its monomeric form contained inactive NF-κB (Fig 7C). Together, these results suggest that the function of human BCL10 as a kinetic determinant of cellular decisions has been retained for over 600 million years of animal evolution.

## Discussion

In this study, we investigated the influence of CBM signalosome assembly on the binary kinetics of transcription factor NF-κB activation, a crucial effector of immune cell function. Using biophysical tools and a single cell reporter of NF-κB activity, we revealed that the all-or-none response of NF-κB has a structural basis in the CBM signalosome, and that it specifically results from a sequence-encoded nucleation barrier in the CARD domain of the adaptor protein, BCL10.

This barrier allows BCL10 to exist in a physiologically supersaturated inactive state that is poised for self-templating polymerization-dependent activation upon stimulus-dependent nucleation by any of four upstream CARD-CC proteins. We characterized in depth one of those -- CARD9 -- and showed that its long coiled coil region drives it into large multimers that facilitate nucleation by lowering the entropic cost to assembling multiple CARD subunits into a nucleating configuration. We reiterate that the role of upstream CARD-CC proteins is primarily kinetic in nature and culminates with the initial nucleating event, as evidenced by our finding that BCL10 continues to polymerize even after the experimental dissolution of CARD nuclei.

Our characterization of the cnidarian CBM proteins suggests that the nucleation barrier is a conserved ancient property of BCL10, and our characterization of hypo- and hyperactive pathogenic mutants of the human proteins further suggests that it is evolutionarily tuned for optimum immune system function, where increases or decreases in the barrier are associated with pathogen susceptibility or cancer, respectively.

Our findings introduce a novel structure-function paradigm. Existing explanations for the preponderance of ordered polymers in immune cell signalosomes have centered on the functions of multivalency at steady state, such as scaffolding and sensitivity enhancement resulting from the cooperativity of homo-oligomerization (Nanson et al., 2019; Vajjhala et al., 2017; Wu and Fuxreiter, 2016). Indeed, many intracellular signaling networks exploit the phase boundaries associated with protein self-association to achieve extraordinary sensitivity to stimulation (Dignon et al., 2020; Gibson et al., 2019; Riback et al., 2017; Yoo et al., 2019). Unlike immune cell signalosomes, however, these are primarily driven by the demixing of disordered proteins from the cellular milieu, and therefore lack nucleation barriers sufficiently large to restrict signaling over biological timescales. That is, liquid-liquid phase separation prevents proteins from achieving a deeply supersaturated soluble state, such as occurs for BCL10. This fact, together with demonstrations that signalosome effectors activate simply through an increase in local proximity independent of adaptor polymer structure (herein, and (Boucher et al., 2018; Shen et al., 2018; Würstle et al., 2010)), suggests that the intricately ordered polymers of CBM and other signalosomes are molecular examples of evolutionary spandrels (Gould and Lewontin, 1979; Manhart and Morozov, 2015), byproducts of evolutionary selection. In this case, evolution has acted *against* alternative, less ordered assemblies that would prematurely desaturate the constituent proteins and preclude the subsequent switch-like activation of effector proteins at the moment in time they are needed. Sequence variants that reduce ordering in BCL10, MAVS, ASC, RIPK3, and potentially other prion-like signalosome polymers would concomitantly reduce the nucleation barrier to their formation, and would therefore be pathogenically ineffective arbiters of inflammation and programmed cell death irrespective of the proteins’ abilities to activate downstream effectors. The only assemblies that have escaped this filter are those whose formation involves a sufficiently large nucleation barrier that they rarely form spontaneously, i.e. highly ordered polymers.

Nevertheless, nucleation is inherently probabilistic and, with enough time, destined to occur even in the absence of stimulation. In embracing supersaturation to execute decisions, immune cells may have also predetermined their fates. That BCL10 and potentially other pro-inflammatory signalosome proteins are constitutively supersaturated implies that cells are thermodynamically predisposed to inflammation, and therefore hints at an ultimate physical basis for the centrality of inflammation in progressive and age-associated diseases (Furman et al., 2019; López-Otín et al., 2013; Taniguchi and Karin, 2018). This fact should guide ongoing efforts to control inflammation for the betterment of human health.

## Acknowledgements

We thank members of the Halfmann lab for critical reading of the manuscript. This work was performed to fulfill, in part, requirements for ARG’s thesis research in the Graduate School of the Stowers Institute for Medical Research. This work was supported by the National Institute Of General Medical Sciences (Award Number R01GM130927, to RH) and the National Institute on Aging (Award Number F99AG068511, to ARG) of the National Institutes of Health, the American Cancer Society (RSG-19-217-01-CCG to RH), and the Stowers Institute for Medical Research. The funders had no role in study design, data collection and analysis, or manuscript preparation. The content is solely the responsibility of the authors and does not necessarily represent the official views of the funders. Original data underlying this manuscript can be accessed from the Stowers Original Data Repository at http://www.stowers.org/research/publications/libpb-1675.

## Materials and methods

### Antibodies and reagents

#### Antibodies used in this study include

Bcl10 Rabbit polyclonal Ab (Cell Signaling Technology, C78F1), CARD9 mouse monoclonal Ab (Santa Cruz Biotechnology, A-8: sc-374569), MALT1 mouse monoclonal Ab (Santa Cruz Biotechnology, B-12: sc-46677). Actin mouse monoclonal Ab (Santa Cruz Biotechnology, C4: sc-47778). mEos3.1 Rabbit polyclonal Ab (Halfmann laboratory, #4530). Alexa-488 Goat anti-Rabbit IgG (Thermo Fisher Scientific, A32731). Anti-mouse IgG Ab HRP (Cell Signaling Technology, 7076), mouse anti-rabbit IgG-HRP (Santa Cruz Biotechnology, sc-2357). Other reagents include: FuGENE HD (promega, E2311), β-glucan peptide (Invivogen, tlrl-bn, ant-zn-05), Hygromycin B (Invivogen, ant-hg-1), Penicillin-Streptomycin (ThermoFisher, 1514014gp), LPS (Sigma-Aldrich, L2880), PMA (BioVision, 1544-5), Dimerizer ligand (AP201870) (Takara, 635058), Halt protease inhibitor (ThermoFisher, 78439), Puromycin (Invivogen, ant-pr-1), Zeocin (Invivoge8).

### Plasmid construction

A list of plasmids used in this study can be found in table S1. Yeast expression plasmids were made as previously described in Khan et al 2018. Briefly, we used a golden gate cloning-compatible vector V08 which contains inverted BsaI sites followed by 4x(EAAAR) linker and mEos3.1. V08 vector drives expression of proteins from a GAL promoter and contains the auxotrophic marker URA3. The m1 mammalian expression vector was constructed in two steps from mEos3.2-N1 (Addgene #54525). First an existing BsaI site in mEos3.2-N1 was removed by introducing a point mutation at nucleotide position (3719) G to A. Second, the existing multiple cloning site in mEos3.2-N1 was replaced via gibson assembly with a fragment containing inverted BsaI sites followed by 4x(EAAAR) to create the golden gate compatible vector, m1. CMV promoter in m1 controls protein expression. m1 was then used to create the m1_C1 vector by removing the inverted BsaI sites upstream of mEos3.1. Then placing a fragment containing 4x(EAAAR) followed by inverted BsaI after mEos3.2 via gibson. MCardinal vector was made by replacing the mEos3.2 in m1 with a synthetic string coding the marker mCardinal via gibson. m16 vector was made by replacing the mEos3.2 in m1 with a synthetic string coding the marker mScarlet-I via gibson. Vector m13 was made by inserting a string containing 3x(DmrB) domains in front of mEos3.1 in m1_C1 and replacing mEos3.2 with mKelly2 marker via gibson. m14 was then made by removing two copies of DmrB using enzyme BamHI. m25 backbone was made by replacing the mCherry sequence in mCherry-CRY2clust (Addgene #105624) with an insert containing inverted BsaI sited followed by 4x(EAAAR) and miRFP670nano.

Lentivirus vectors were made as follows. The lentivirus vector containing the NF-κB reporter, phage_NFkB-TSapp, was made by replacing the luciferase sequences with the fluorescent protein T-Sapphire in pHAGE NFKB-TA-LUC-UBC-dTomato-W (Addgene #49335). Additionally we replaced the marker tdTomato in Addgene #49335 with the resistance cassette for HygromycinB. Lentivirus vector for expression of EYFP-p65, pLV_EYFP-p65, was made by replacing the HygromycinB resistance in pLV-EF1a-IRES-Hygro (Addgene #85134) with Zeocin resistance cassette and inserting the p65 coding sequence before the IRES site. To create lentiviral vectors for the expression of CARD9 phused with miRFP670 and Cry2, we inserted via gibson the coding sequence of cloned inserts in m25 into pLV-EF1a-IRES-Hygro. Finally, for the doxycycline controlled lentiviral vectors we cloned the respective coding sequences from *BCL10* cloned in m1 or m16 vectors into pCW57.1 (Addgene #41393).

Inserts were ordered as GeneArt Strings (Thermo Fisher) flanked by Type IIs restriction sites for ligation between BsaI sites in V08, V12, m1, and other vectors derived from m1. All other inserts were cloned into respective vectors via gibson assembly between the promoter and respective fluorescent or tag marker. All plasmids were verified by Sanger sequencing.

### Yeast strain construction

For DAmFRET experiments we employed strain rhy1713 which was made as previously described in Khan et al 2018. To create artificial intracellular seeds of CARD9 variants, CARD10, CARD11 and CARD14, sequences were fused to a constitutive condensate-forming protein, μNS (471–721), hereafter “μNS”. Yeast strain rhy2153 was created by replacing the HO locus in rhy1734 with a cassette consisting of: natMX followed by the tetO7 promoter followed by counterselectable URA3 ORFs derived from C. albicans and K. lactis, followed by μNS-mCardinal. To create strains expressing the fusion CARD9-mCardinal-μNS and others, AseI digests of yeast expression plasmids containing desired proteins were transformed into rhy2153 to replace the counterselectable URA3 ORFs with the gene of interest. The resulting strains express the proteins of interest fused to μNS-mCardinal, under the control of a doxycycline-repressible promoter. Transformants were selected for 5-FOA resistance and validated by PCR. See Table S2 for a list of all strains used in this study.

For measuring nucleating interactions diploid strains were maintained on doxycycline (40 mg/ml) until initial culture for DAmFRET assay.

### Cell culture

HEK293T cells and THP-1 cells were purchased from ATCC. HEK293T cells were grown in Dulbecco’s Modified Eagle’s Medium (DMEM) with L-glutamine, 10% fetal bovine serum (FBS) and PenStrep 100U/mL. THP-1 cells were grown in Roswell Park Memorial Institute (RPMI) medium 1640 with L-glutamine and 10% FBS. All cells were grown at 37°C in a 5% CO2 atmosphere incubator. Cell lines were regularly tested for mycoplasma using the Universal mycoplasma detection kit (ATCC, #30-1012K).

### Transient transfections

HEK293T cells were plated in 6-well culture plates at 8.0 × 10^5^ cells/well in DMEM. The next day, 2.0 ug DNA of expression plasmids were mixed 150 μL Opti-MEM and transfected using FuGENE HD (Promega) at a DNA to FuGENE HD ratio of (1:3). Cells were treated with either PMA or Dimerizer ligand at described concentrations 24 hours after transfection. Cells were harvested for flow cytometry analysis after a total of 48 hours. For microscopy experiments, cells were plated directly into 35mm glass-bottom dishes (iBidi, μ-Dish 35 mm, high Glass Bottom) at a concentration of 0.8 × 10^5^ cells/dish and transfected as previously mentioned.

### Generation of stable cell lines

For generation of stable cell lines, constructs were packaged into lentivirus in a 10 cm plate 60% confluent of HEK293T cells using the TransIT-LT1 (Mirus Bio, MIR2300) transfection reagent and 7 μg of the vector, 7 μg psPAX2 and 1 μg pVSV-G. After 48 hours, media was collected and centrifuged at 3000g to remove cell debris. At this point, media containing lentivirus was used or stored at −80°C. For HEK293T adherent cell transduction, cells were plated in 6-well culture plates at 50% density. Next day, media was replaced with media containing lentivirus at a multiplicity of infection of 1, supplemented with Polybrene 5 ug/ml (Sigma Aldrich, TR-1003-G). For THP-1 suspension cell transduction, cells were spinfected with virus containing media for 1 h at 1,000g at room temperature supplemented with Polybrene 5 μg/ml. For transduction of phageNF-κB-TSapp, THP-1 and HEK293T cells were selected with Hygromycin B (350 mg/mL) for 14 days and used for further analysis or complementary transductions. For transduction of pCW57.1_Bcl10-mScarlet, pwtBcl10-mScarlet, HEK293T cells were selected with Puromycin (1 ug/mL) for 7 days. After this time, cells were sorted for positive expression of mScarlet and expanded in continuing selection with puromycin. For transduction of expression clones tagged with mEos3.1, THP-1 cells were selected with Puromycin (1 ug/mL) for 7 days. Cells were sorted for positive expression of mEos3.1 and expanded for further experiments with continued selection in puromycin. For transduction of pLV_EYFP-p65, HEK293T *BCL10*-KO cells reconstituted with pBcl10-mScarlet, were transduced and selected with Zeocin (300 ug/mL). Positive cells expressing EYFP and mScarlet were sorted and expanded in selection media containing drugs. Plasmid pCARD9-CARD_miRFP-Cry2 was transduced into HEK293T cells. Cells were selected with Hygromycin B (350 ug/mL) for 7 days and then sorted for positive expression of miRFP670nano.

### CRISPR-Cas9 mediated knock-out

To generate HEK293T *BCL10*-KO and MALT KO cell line, cells were nucleofected (Neon Transfection system, Thremo) with a ribonucleoprotein (RNP) mix of sgRNA for *BCL10* exon1 or MALT1 exon1 and Cas9 protein. After this, cells were pooled and submitted for targeted deep sequencing to find the insertion/deletion (InDel) ratio. Subsequently, cells were prepared for single-cell sorting into four 96-well plates and expanded to obtain a duplicate for each well. At this stage cells were lysed and submitted for deep sequencing to find the genome edited cells. Selected wells were further expanded to validate via protein immuno detection. To generate THP-1 *BCL10*-KO cells, sgRNA targeting *BCL10* exon1 was cloned into the lentiCRISPR v2-Blast (Addgene #83480). This vector was packaged into lentivirus as previously described. THP-1 cells were transduced via spinfection with lentivirus and supplemented with polybrene. 24 hours after spinfection, media was replaced. 48 hours after spinfection, cells were selected with Blasticidin (1 ug/ml). After 10 days of blasticidin selection, single-cell clonal expansion was done by serial dilution of resistant cells to achieve complete knockouts. Selected wells were analyzed by immuno detection to confirm the absence of BCL10 protein.

### Yeast preparation for DAmFRET

We performed DAmFRET analysis as previously described in Khan et al. 2018. Expression plasmids were transformed into specified strains following a standard lithium acetate protocol (see table X for list of expression plasmids). Yeast transformants were selected in SD-URA plates. Briefly, single transformant colonies were inoculated in 200 μl of SD-URA in a microplate well and incubated in a Heidolph Titramax platform shaker at 30°C, 1350 RPM overnight without presence of Dox. Cells were washed with sterile water, resuspended in galactose-containing media, and allowed to continue incubating for approximately 16 h. Microplates were then illuminated for 25 min with 320–500 nm violet light to photoconvert a fraction of mEos3.1 molecules from a green (516 nm) form to a red form (581 nm). At this point cells were either used to collect microscopy data or continue the DAmFRET protocol.

### Mammalian cell preparation for DAmFRET

We performed DAmFRET in HEK293T cells as previously described in Venkatesan et al. Briefly, cells were transfected as previously described. 48 hours after transfection, cells were trypsinized and washed in PBS supplemented with 10 mM EDTA. Then, samples were fixed in 4% paraformaldehyde for 5 min with constant shaking. Cells were washed two additional times with PBS + EDTA and transferred to a 96-well plate. The sample-containing plate was photoconverted as previously described for yeast samples. After photoconversion, samples were analysed in a ZE5 cell analyzer cytometer.

### DAmFRET cytometric data collection

DAmFRET data was collected on a ZE5 cell analyzer cytometer or an ImageStreamx MkII imaging cytometer (Amnis) at x60 magnification as previously described in Khan et al. Autofluorescence was detected with 405 nm excitation and 460/22 nm emission; SSC and FSC were detected with 488 nm excitation and 488/10 nm emission. Donor and FRET fluorescence were detected with 488 nm excitation and 425/35 nm or 593/52 nm emission, respectively. Acceptor fluorescence was excited with 561 nm excitation and 589/15 nm emission. Approximately 500,000 events were collected per sample. Data compensation was done in the built-in tool for compensation (Everest software V1.1) on single-color controls: non-photoconverted mEos3.1 and dsRed2 (as a proxy for the red form of mEos3.1). For nucleating interactions, all regular channels for DAmFRET were collected with the addition of collection of mCardinal intensity by 561 nm excitation and 670/30 nm emission.

DAmFRET coupled with the NF-κB reporter data was collected as previously mentioned with the following modifications. HEK293T cells were treated as previously described in the DAmFRET protocol. In addition to the regular channel for DAmFRET data acquisition, T-Sapphire expression was detected using 405 nm excitation and 525/50 nm emission.

### DAmFRET analysis

Data was processed on FCS Express Plus 6.04.0015 software (De Novo). Events were gated for single unbudded cells by FSC vs. SSC, followed by gating of live cells with low autofluorescence and donor positive. Live gate was then selected for double positives (donor and acceptor). Plots represent the distribution of AmFRET (FRET intensity/acceptor intensity) vs. Acceptor intensity (protein expression). For DAmFRET data in mammalian cells, cells were first gated for single cells followed by gating double positive expressing cells donor, acceptor). In the cases for DAmFRET coupled to NF-κB reporter, an additional gating was performed on the DAmFRET plot to quantify the intensity of T-Sapphire.

Weibull fit analysis was performed as previously described in Khan et al 2018. Briefly, a negative DAmFRET histogram was defined and used to set a negative gating strategy. DAmFRET plots were divided into 64 logarithmically spaced bins and then determined the fraction of cells that have AmFRET values above the defined negative control. We applied this approach to all DAmFRET plots. This analysis provides a gross percentage of the fraction of cells containing assembled protein. The data of fraction of assembled cells in each bin was fit to a Weibull distribution to extract the statistics EC50 and delta.

### Fluorescence microscopy

Imaging of yeast and mammalian cells was done in an LSM 780 microscope with a 63x Plan-Apochromat (NA =1.40) objective. T-Sapphire excitation was done with a 405nm laser. mEos3.1 and mScarlet-I excitation was made with a 488 nm and 561 nm laser respectively. To quantify the percentage of cells containing polymers, images were subjected to an in-house Fiji (https://imagej.net/software/fiji/) adapted implementation of Cellpose (https://www.nature.com/articles/s41592-020-01018-x, https://www.cellpose.org/) analysis for cellular segmentation for both whole cell contour and nucleus. The Cellpose generated regions of interest (ROIs) were used to measure specified imaging channels. For each experiment more than 200 cells were quantified in an unbiased way. Cells were defined as positive for containing polymers if the Coefficient of variation (CV, standard deviation divided by the mean intensity) for the fluorescent channel was above 0.8 as defined by the distribution of values.

For time lapse imaging, fluorescence images were taken using a spinning-disk confocal microscope (Nikon, CSU-W1) with a 60X Plan Apochromat objective (NA=1.40) and a Flash 4 sCMOS camera (Hamamatsu). Samples were maintained at 37 °C and 5% CO2 with a stage top incubator (Okolab). Prior to stimulation, cells were treated with Hoechst 33342 (0.5 ug/ml) for 30 min. 405 nm laser at 20% was used to collect emission of Hoechst for 50 ms per frame. 514 nm and 561 nm laser were used at 25% to collect EYFP and m-Scarlet emission for 300ms, respectively. Images were captured every 10 minutes for p65 nuclear translocation experiments. ROIs were generated by Cellpose segmentation algorithm around each cell contour and nucleus on the EYFP and Hoechst image respectively. These ROIs were then used to measure the area, mean, standard deviation, and integrated density of each cell on the EYFP and mScarlet fluorescence channels. For each cell, we computed the nuclear to cytosol ratio of integrated density for the EYFP channel. The same ROIs were used to compute the coefficient of variation (CV, standard deviation divided by the mean intensity) of the mScarlet-I fluorescence.

### Optogenetic activation

Cry2clust optogenetic experiments were made in an LSM 780 microscope with a 63x Plan-Apochromat (NA = 1.40) objective in the integration mode. Samples were maintained at 37 °C and 5% CO2 with a microscope incubator. To stimulate Cry2clust we used the 488 nm laser at a power setting of 50% for a pulse of 1 second, which is the amount of time it took to scan the user generated region of interest, unless indicated otherwise. 561 and 633 nm lasers were used for imaging mScarlet-I and miRFP670nano, respectively. To track BCL10 nucleation dependent on CARD9-Cry2 we imaged cells before and after stimulation every 2 minutes. For the experiments to quantify the nuclear translocation of p65 to the nucleus, EYFP was imaged using the 514 nm laser. Imaging EYFP with the 514 laser resulted in undesired Cry2clust restimulation. To avoid additional Cry2clust stimulation in our experiments, EYFP was only imaged at the beginning of the experiment and in the last frame. We computed the nuclear to cytosol ratio of EYFP as previously described. Similarly, we calculated the CV for mScarlet-I intensity as mentioned above.

### Protein immunodetection

The Wes platform (ProteinSimple) was used to perform capillary based protein immunodetection. We followed the recommended protocol from the manufacturer. Briefly, cell lysates were diluted with 0.1 × Sample Buffer. Subsequently, 4 parts of diluted sample were combined with 1 part 5 × Fluorescent Master Mix (containing 5 × sample buffer, 5 × fluorescent standard, and 200 mM DTT) then boiled at 95 °C for 5 min. Our sample final protein concentration for each capillary was 1 ug/ml. After the denaturing step, an assay plate was filled with samples, blocking reagent, primary antibodies (1:50 dilution for mEos3.1, MALT1 and actin, 1:10 dilution for BCL10, and 1:150 dilution for CARD9), HRP-conjugated secondary antibodies and chemiluminescent substrate. A biotinylated ladder provided molecular weight standards for each assay. Once the assay plate was set up, electrophoretic protein separation and immunodetection was carried out in the fully automated capillary system. Data was processed using the open source software Compass to extract the intensities for the peaks corresponding to the expected molecular weight of proteins of interest.

### Quantification and statistical analysis

For two sample comparisons, two-sided Student’s t-tests were used for significance testing unless stated otherwise. The graphs represent the means ± SEM of independent biological experiments unless stated otherwise. Statistical analysis was performed using GraphPad Prism 9 and R packages.

## References

Boucher D, Monteleone M, Coll RC, Chen KW, Ross CM, Teo JL, Gomez GA, Holley CL, Bierschenk D, Stacey KJ, Yap AS, Bezbradica JS, Schroder K. 2018. Caspase-1 self-cleavage is an intrinsic mechanism to terminate inflammasome activity. J Exp Med 215:827–840. doi:10.1084/jem.20172222

Cai X, Chen J, Xu H, Liu S, Jiang Q-X, Halfmann R, Chen ZJ. 2014. Prion-like polymerization underlies signal transduction in antiviral immune defense and inflammasome activation. Cell 156:1207–1222. doi:10.1016/j.cell.2014.01.063

Compagno M, Lim WK, Grunn A, Nandula SV, Brahmachary M, Shen Q, Bertoni F, Ponzoni M, Scandurra M, Califano A, Bhagat G, Chadburn A, Dalla-Favera R, Pasqualucci L. 2009. Mutations of multiple genes cause deregulation of NF-kappaB in diffuse large B-cell lymphoma. Nature 459:717–721. doi:10.1038/nature07968

David L, Li Y, Ma J, Garner E, Zhang X, Wu H. 2018. Assembly mechanism of the CARMA1-BCL10-MALT1-TRAF6 signalosome. Proc Natl Acad Sci USA 115:1499–1504. doi:10.1073/pnas.1721967115

Demarco B, Grayczyk JP, Bjanes E, Le Roy D, Tonnus W, Assenmacher C-A, Radaelli E, Fettrelet T, Mack V, Linkermann A, Roger T, Brodsky IE, Chen KW, Broz P. 2020. Caspase-8-dependent gasdermin D cleavage promotes antimicrobial defense but confers susceptibility to TNF-induced lethality. Sci Adv 6. doi:10.1126/sciadv.abc3465

Dignon GL, Best RB, Mittal J. 2020. Biomolecular phase separation: from molecular driving forces to macroscopic properties. Annu Rev Phys Chem 71:53–75. doi:10.1146/annurev-physchem-071819-113553

Franklin BS, Bossaller L, De Nardo D, Ratter JM, Stutz A, Engels G, Brenker C, Nordhoff M, Mirandola SR, Al-Amoudi A, Mangan MS, Zimmer S, Monks BG, Fricke M, Schmidt RE, Espevik T, Jones B, Jarnicki AG, Hansbro PM, Busto P, Latz E. 2014. The adaptor ASC has extracellular and “prionoid” activities that propagate inflammation. Nat Immunol 15:727–737. doi:10.1038/ni.2913

Furman D, Campisi J, Verdin E, Carrera-Bastos P, Targ S, Franceschi C, Ferrucci L, Gilroy DW, Fasano A, Miller GW, Miller AH, Mantovani A, Weyand CM, Barzilai N, Goronzy JJ, Rando TA, Effros RB, Lucia A, Kleinstreuer N, Slavich GM. 2019. Chronic inflammation in the etiology of disease across the life span. Nat Med 25:1822–1832. doi:10.1038/s41591-019-0675-0

Gehring T, Seeholzer T, Krappmann D. 2018. BCL10 - Bridging CARDs to Immune Activation. Front Immunol 9:1539. doi:10.3389/fimmu.2018.01539

Gibson BA, Doolittle LK, Schneider MWG, Jensen LE, Gamarra N, Henry L, Gerlich DW, Redding S, Rosen MK. 2019. Organization of chromatin by intrinsic and regulated phase separation. Cell 179:470–484.e21. doi:10.1016/j.cell.2019.08.037

Ginster S, Bardet M, Unterreiner A, Malinverni C, Renner F, Lam S, Freuler F, Gerrits B, Voshol J, Calzascia T, Régnier CH, Renatus M, Nikolay R, Israël L, Bornancin F. 2017. Two Antagonistic MALT1 Auto-Cleavage Mechanisms Reveal a Role for TRAF6 to Unleash MALT1 Activation. PLoS ONE 12:e0169026. doi:10.1371/journal.pone.0169026

Glocker E-O, Hennigs A, Nabavi M, Schäffer AA, Woellner C, Salzer U, Pfeifer D, Veelken H, Warnatz K, Tahami F, Jamal S, Manguiat A, Rezaei N, Amirzargar AA, Plebani A, Hannesschläger N, Gross O, Ruland J, Grimbacher B. 2009. A homozygous CARD9 mutation in a family with susceptibility to fungal infections. N Engl J Med 361: 1727–1735. doi:10.1056/NEJMoa0810719

Gould SJ, Lewontin RC. 1979. The spandrels of San Marco and the Panglossian paradigm: a critique of the adaptationist programme. Proc R Soc Lond, B, Biol Sci 205:581–598. doi:10.1098/rspb.1979.0086

Gross O, Gewies A, Finger K, Schäfer M, Sparwasser T, Peschel C, Förster I, Ruland J. 2006. Card9 controls a non-TLR signalling pathway for innate anti-fungal immunity. Nature 442:651–656. doi:10.1038/nature04926

Holliday MJ, Witt A, Rodríguez Gama A, Walters BT, Arthur CP, Halfmann R, Rohou A, Dueber EC, Fairbrother WJ. 2019. Structures of autoinhibited and polymerized forms of CARD9 reveal mechanisms of CARD9 and CARD11 activation. Nat Commun 10:3070. doi:10.1038/s41467-019-10953-z

Hou F, Sun L, Zheng H, Skaug B, Jiang Q-X, Chen ZJ. 2011. MAVS forms functional prion-like aggregates to activate and propagate antiviral innate immune response. Cell 146:448–461. doi:10.1016/j.cell.2011.06.041

Howes A, O’Sullivan PA, Breyer F, Ghose A, Cao L, Krappmann D, Bowcock AM, Ley SC. 2016. Psoriasis mutations disrupt CARD14 autoinhibition promoting BCL10-MALT1-dependent NF-κB activation. Biochem J 473:1759–1768. doi:10.1042/BCJ20160270

Hsu Y-MS, Zhang Y, You Y, Wang D, Li H, Duramad O, Qin X-F, Dong C, Lin X. 2007. The adaptor protein CARD9 is required for innate immune responses to intracellular pathogens. Nat Immunol 8:198–205. doi:10.1038/ni1426

Jordan CT, Cao L, Roberson EDO, Pierson KC, Yang C-F, Joyce CE, Ryan C, Duan S, Helms CA, Liu Y, Chen Y, McBride AA, Hwu W-L, Wu J-Y, Chen Y-T, Menter A, Goldbach-Mansky R, Lowes MA, Bowcock AM. 2012. PSORS2 is due to mutations in CARD14. Am J Hum Genet 90:784–795. doi:10.1016/j.ajhg.2012.03.012

Jumper J, Evans R, Pritzel A, Green T, Figurnov M, Ronneberger O, Tunyasuvunakool K, Bates R, Žídek A, Potapenko A, Bridgland A, Meyer C, Kohl SAA, Ballard AJ, Cowie A, Romera-Paredes B, Nikolov S, Jain R, Adler J, Back T, Hassabis D. 2021. Highly accurate protein structure prediction with AlphaFold. Nature 596:583–589. doi:10.1038/s41586-021-03819-2

Kandola T, Zhang J, Venkatesan S, Lerbakken B, Blanck JF, Wu J, Unruh J, Berry P, Lange JL, Von Schulze A, Box A, Cook M, Sagui C, Halfmann R. 2021. The polyglutamine amyloid nucleus in living cells is monomeric and has competing dimensions of order. BioRxiv. doi:10.1101/2021.08.29.458132

Kellogg RA, Tian C, Lipniacki T, Quake SR, Tay S. 2015. Digital signaling decouples activation probability and population heterogeneity. eLife 4:e08931. doi:10.7554/eLife.08931

Khan T, Kandola TS, Wu J, Venkatesan S, Ketter E, Lange JJ, Rodríguez Gama A, Box A, Unruh JR, Cook M, Halfmann R. 2018. Quantifying nucleation in vivo reveals the physical basis of prion-like phase behavior. Mol Cell 71:155–168.e7. doi:10.1016/j.molcel.2018.06.016

Kingeter LM, Paul S, Maynard SK, Cartwright NG, Schaefer BC. 2010. Cutting edge: TCR ligation triggers digital activation of NF-kappaB. J Immunol 185:4520–4524. doi:10.4049/jimmunol.1001051

Koch D. 2020. Homo-Oligomerisation in Signal Transduction: Dynamics, Homeostasis, Ultrasensitivity, Bistability. J Theor Biol 499:110305. doi:10.1016/j.jtbi.2020.110305

Kuper-Hommel MJJ, Schreuder MI, Gemmink AH, van Krieken JHJM. 2013. T(14;18)(q32;q21) involving MALT1 and IGH genes occurs in extranodal diffuse large B-cell lymphomas of the breast and testis. Mod Pathol 26:421–427. doi:10.1038/modpathol.2012.170

Lanternier F, Mahdaviani SA, Barbati E, Chaussade H, Koumar Y, Levy R, Denis B, Brunel A-S, Martin S, Loop M, Peeters J, de Selys A, Vanclaire J, Vermylen C, Nassogne M-C, Chatzis O, Liu L, Migaud M, Pedergnana V, Desoubeaux G, Puel A. 2015. Inherited CARD9 deficiency in otherwise healthy children and adults with Candida species-induced meningoencephalitis, colitis, or both. J Allergy Clin Immunol 135:1558–68.e2. doi:10.1016/j.jaci.2014.12.1930

Lanternier F, Pathan S, Vincent QB, Liu L, Cypowyj S, Prando C, Migaud M, Taibi L, Ammar-Khodja A, Stambouli OB, Guellil B, Jacobs F, Goffard J-C, Schepers K, Del Marmol V, Boussofara L, Denguezli M, Larif M, Bachelez H, Michel L, Puel A. 2013. Deep dermatophytosis and inherited CARD9 deficiency. N Engl J Med 369:1704–1714. doi:10.1056/NEJMoa1208487

Latty SL, Sakai J, Hopkins L, Verstak B, Paramo T, Berglund NA, Cammarota E, Cicuta P, Gay NJ, Bond PJ, Klenerman D, Bryant CE. 2018. Activation of Toll-like receptors nucleates assembly of the MyDDosome signaling hub. eLife 7. doi:10.7554/eLife.31377

Lenz G, Davis RE, Ngo VN, Lam L, George TC, Wright GW, Dave SS, Zhao H, Xu W, Rosenwald A, Ott G, Muller-Hermelink HK, Gascoyne RD, Connors JM, Rimsza LM, Campo E, Jaffe ES, Delabie J, Smeland EB, Fisher RI, Staudt LM. 2008. Oncogenic CARD11 mutations in human diffuse large B cell lymphoma. Science 319:1676–1679. doi:10.1126/science.1153629

Liu T, Yamaguchi Y, Shirasaki Y, Shikada K, Yamagishi M, Hoshino K, Kaisho T, Takemoto K, Suzuki T, Kuranaga E, Ohara O, Miura M. 2014. Single-cell imaging of caspase-1 dynamics reveals an all-or-none inflammasome signaling response. Cell Rep 8:974–982. doi:10.1016/j.celrep.2014.07.012

López-Otín C, Blasco MA, Partridge L, Serrano M, Kroemer G. 2013. The hallmarks of aging. Cell 153:1194–1217. doi:10.1016/j.cell.2013.05.039

Lu A, Magupalli VG, Ruan J, Yin Q, Atianand MK, Vos MR, Schröder GF, Fitzgerald KA, Wu H, Egelman EH. 2014. Unified polymerization mechanism for the assembly of ASC-dependent inflammasomes. Cell 156:1193–1206. doi:10.1016/j.cell.2014.02.008

Manhart M, Morozov AV. 2015. Protein folding and binding can emerge as evolutionary spandrels through structural coupling. Proc Natl Acad Sci USA 112:1797–1802. doi:10.1073/pnas.1415895112

Matyszewski M, Morrone SR, Sohn J. 2018. Digital signaling network drives the assembly of the AIM2-ASC inflammasome. Proc Natl Acad Sci USA 115:E1963–E1972. doi:10.1073/pnas.1712860115

Mompeán M, Li W, Li J, Laage S, Siemer AB, Bozkurt G, Wu H, McDermott AE. 2018. The structure of the necrosome RIPK1-RIPK3 core, a human hetero-amyloid signaling complex. Cell 173:1244–1253.e10. doi:10.1016/j.cell.2018.03.032

Muñoz JF, Delorey T, Ford CB, Li BY, Thompson DA, Rao RP, Cuomo CA. 2019. Coordinated host-pathogen transcriptional dynamics revealed using sorted subpopulations and single macrophages infected with Candida albicans. Nat Commun 10:1607. doi:10.1038/s41467-019-09599-8

Nanson JD, Kobe B, Ve T. 2019. Death, TIR, and RHIM: Self-assembling domains involved in innate immunity and cell-death signaling. J Leukoc Biol 105:363–375. doi:10.1002/JLB.MR0318-123R

O’Carroll A, Coyle J, Gambin Y. 2020. Prions and Prion-like assemblies in neurodegeneration and immunity: The emergence of universal mechanisms across health and disease. Semin Cell Dev Biol 99:115–130. doi:10.1016/j.semcdb.2019.11.012

Park H, Kim NY, Lee S, Kim N, Kim J, Heo WD. 2017. Optogenetic protein clustering through fluorescent protein tagging and extension of CRY2. Nat Commun 8:30. doi:10.1038/s41467-017-00060-2

Posey AE, Ruff KM, Lalmansingh JM, Kandola TS, Lange JJ, Halfmann R, Pappu RV. 2021. Mechanistic inferences from analysis of measurements of protein phase transitions in live cells. J Mol Biol 433:166848. doi:10.1016/j.jmb.2021.166848

Qiao Q, Yang C, Zheng C, Fontán L, David L, Yu X, Bracken C, Rosen M, Melnick A, Egelman EH, Wu H. 2013. Structural architecture of the CARMA1/Bcl10/MALT1 signalosome: nucleation-induced filamentous assembly. Mol Cell 51:766–779. doi:10.1016/j.molcel.2013.08.032

Riback JA, Katanski CD, Kear-Scott JL, Pilipenko EV, Rojek AE, Sosnick TR, Drummond DA. 2017. Stress-Triggered Phase Separation Is an Adaptive, Evolutionarily Tuned Response. Cell 168:1028–1040.e19. doi:10.1016/j.cell.2017.02.027

Rodríguez Gama A, Miller T, Halfmann R. 2021. Mechanics of a molecular mousetrap-nucleation-limited innate immune signaling. Biophys J 120:1150–1160. doi:10.1016/j.bpj.2021.01.007

Ruland J, Duncan GS, Elia A, del Barco Barrantes I, Nguyen L, Plyte S, Millar DG, Bouchard D, Wakeham A, Ohashi PS, Mak TW. 2001. Bcl10 is a positive regulator of antigen receptor-induced activation of NF-kappaB and neural tube closure. Cell 104:33–42. doi:10.1016/s0092-8674(01)00189-1

Ruland J, Hartjes L. 2019. CARD-BCL-10-MALT1 signalling in protective and pathological immunity. Nat Rev Immunol 19:118–134. doi:10.1038/s41577-018-0087-2

Schmitz AM, Morrison MF, Agunwamba AO, Nibert ML, Lesser CF. 2009. Protein interaction platforms: visualization of interacting proteins in yeast. Nat Methods 6:500–502. doi:10.1038/nmeth.1337

Shen C, Pei J, Guo X, Zhou L, Li Q, Quan J. 2018. Structural basis for dimerization of the death effector domain of the F122A mutant of Caspase-8. Sci Rep 8:16723. doi:10.1038/s41598-018-35153-5

Sommer K, Guo B, Pomerantz JL, Bandaranayake AD, Moreno-García ME, Ovechkina YL, Rawlings DJ. 2005. Phosphorylation of the CARMA1 linker controls NF-kappaB activation. Immunity 23:561–574. doi:10.1016/j.immuni.2005.09.014

Staal J, Driege Y, Haegman M, Borghi A, Hulpiau P, Lievens L, Gul IS, Sundararaman S, Gonçalves A, Dhondt I, Pinzón JH, Braeckman BP, Technau U, Saeys Y, van Roy F, Beyaert R. 2018. Ancient Origin of the CARD-Coiled Coil/Bcl10/MALT1-Like Paracaspase Signaling Complex Indicates Unknown Critical Functions. Front Immunol 9:1136. doi:10.3389/fimmu.2018.01136

Strasser D, Neumann K, Bergmann H, Marakalala MJ, Guler R, Rojowska A, Hopfner K-P, Brombacher F, Urlaub H, Baier G, Brown GD, Leitges M, Ruland J. 2012. Syk kinase-coupled C-type lectin receptors engage protein kinase C-σ to elicit Card9 adaptor-mediated innate immunity. Immunity 36:32–42. doi:10.1016/j.immuni.2011.11.015

Taniguchi K, Karin M. 2018. NF-κB, inflammation, immunity and cancer: coming of age. Nat Rev Immunol 18:309–324. doi:10.1038/nri.2017.142

Tay S, Hughey JJ, Lee TK, Lipniacki T, Quake SR, Covert MW. 2010. Single-cell NF-kappaB dynamics reveal digital activation and analogue information processing. Nature 466:267–271. doi:10.1038/nature09145

Vajjhala PR, Ve T, Bentham A, Stacey KJ, Kobe B. 2017. The molecular mechanisms of signaling by cooperative assembly formation in innate immunity pathways. Mol Immunol 86:23–37. doi:10.1016/j.molimm.2017.02.012

Venkatesan S, Kandola TS, Rodríguez-Gama A, Box A, Halfmann R. 2019. Detecting and characterizing protein self-assembly in vivo by flow cytometry. J Vis Exp. doi:10.3791/59577

Wang D, You Y, Case SM, McAllister-Lucas LM, Wang L, DiStefano PS, Nuñez G, Bertin J, Lin X. 2002. A requirement for CARMA1 in TCR-induced NF-kappa B activation. Nat Immunol 3:830–835. doi:10.1038/ni824

Willis TG, Jadayel DM, Du MQ, Peng H, Perry AR, Abdul-Rauf M, Price H, Karran L, Majekodunmi O, Wlodarska I, Pan L, Crook T, Hamoudi R, Isaacson PG, Dyer MJ. 1999. Bcl10 is involved in t(1;14)(p22;q32) of MALT B cell lymphoma and mutated in multiple tumor types. Cell 96:35–45. doi:10.1016/s0092-8674(00)80957-5

Wilson AA, Kwok LW, Porter EL, Payne JG, McElroy GS, Ohle SJ, Greenhill SR, Blahna MT, Yamamoto K, Jean JC, Mizgerd JP, Kotton DN. 2013. Lentiviral delivery of RNAi for in vivo lineage-specific modulation of gene expression in mouse lung macrophages. Mol Ther 21:825–833. doi:10.1038/mt.2013.19

Wu H, Fuxreiter M. 2016. The Structure and Dynamics of Higher-Order Assemblies: Amyloids, Signalosomes, and Granules. Cell 165:1055–1066. doi:10.1016/j.cell.2016.05.004

Würstle ML, Laussmann MA, Rehm M. 2010. The caspase-8 dimerization/dissociation balance is a highly potent regulator of caspase-8,-3,-6 signaling. J Biol Chem 285:33209–33218. doi:10.1074/jbc.M110.113860

Yan B, Chen G, Saigal K, Yang X, Jensen ST, Van Waes C, Stoeckert CJ, Chen Z. 2008. Systems biology-defined NF-kappaB regulons, interacting signal pathways and networks are implicated in the malignant phenotype of head and neck cancer cell lines differing in p53 status. Genome Biol 9:R53. doi:10.1186/gb-2008-9-3-r53

Yoo H, Triandafillou C, Drummond DA. 2019. Cellular sensing by phase separation: Using the process, not just the products. J Biol Chem 294:7151–7159. doi:10.1074/jbc.TM118.001191

Zhang Q, Siebert R, Yan M, Hinzmann B, Cui X, Xue L, Rakestraw KM, Naeve CW, Beckmann G, Weisenburger DD, Sanger WG, Nowotny H, Vesely M, Callet-Bauchu E, Salles G, Dixit VM, Rosenthal A, Schlegelberger B, Morris SW. 1999. Inactivating mutations and overexpression of BCL10, a caspase recruitment domain-containing gene, in MALT lymphoma with t(1;14)(p22;q32). Nat Genet 22:63–68. doi:10.1038/8767

Zhong X, Chen B, Yang L, Yang Z. 2018. Molecular and physiological roles of the adaptor protein CARD9 in immunity. Cell Death Dis 9:52. doi:10.1038/s41419-017-0084-6

Zhou T, Souzeau E, Sharma S, Siggs OM, Goldberg I, Healey PR, Graham S, Hewitt AW, Mackey DA, Casson RJ, Landers J, Mills R, Ellis J, Leo P, Brown MA, MacGregor S, Burdon KP, Craig JE. 2016. Rare variants in optic disc area gene CARD10 enriched in primary open-angle glaucoma. Mol Genet Genomic Med 4:624–633. doi:10.1002/mgg3.248

