## Supplemental Figures for "A nucleation barrier spring-loads the CBM signalosome for binary activation"

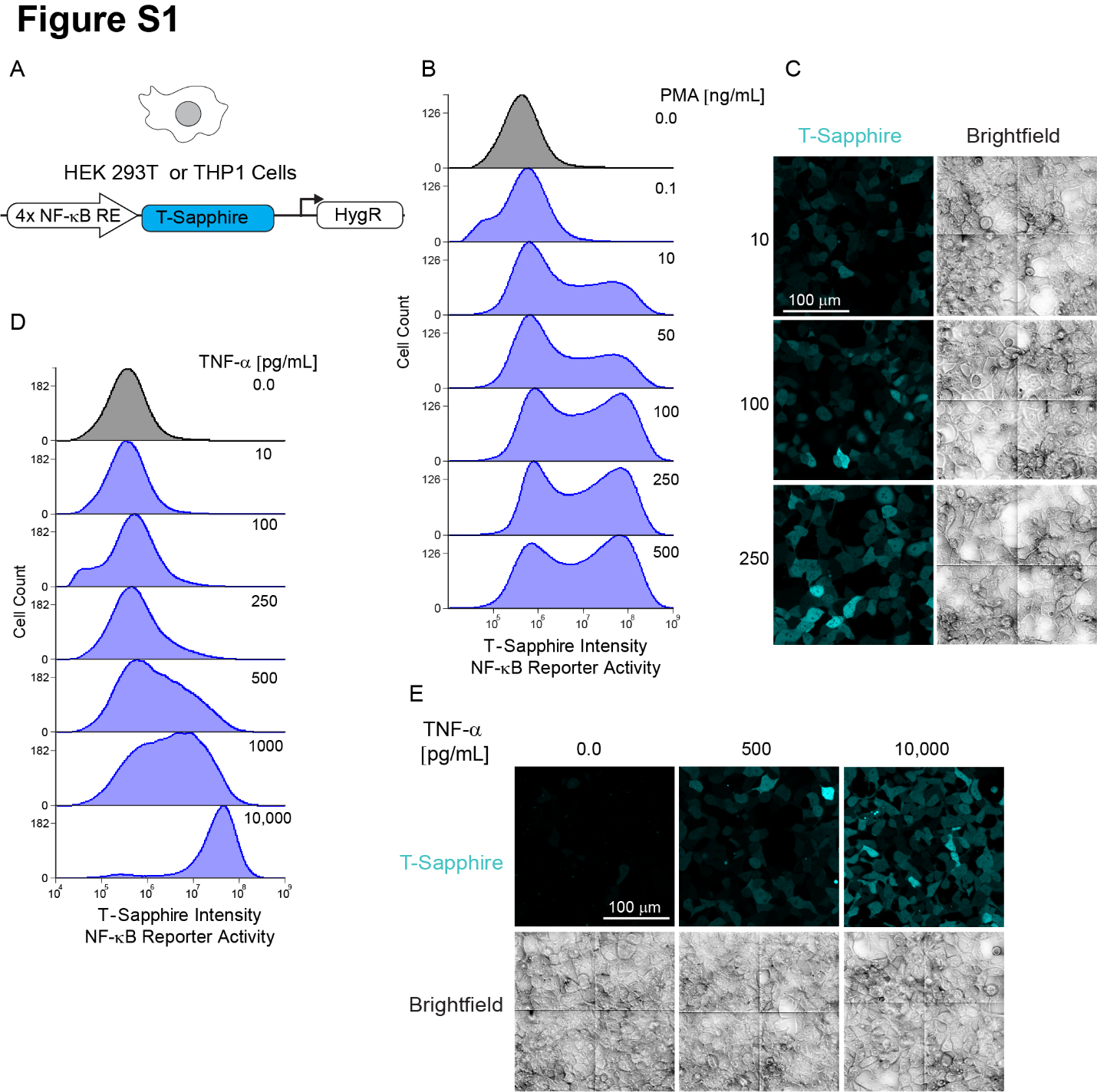

Supplementary Figure 1.

(A) Schematic of the NF-κB reporter integrated into 293T and THP-1 cell lines. The reporter contains 4 minimal response elements of the NF-κB transcription factor followed by the coding sequence of the fluorescence protein T-Sapphire. Cells were selected for hygromycin resistance after lentiviral transduction.

(B) Histograms of T-Sapphire expression for 293T WT cells containing the NF-κB reporter after 24 hours of stimulation with the indicated doses of PMA.

(C) Microscopy images of 293T cells containing the NF-κB reporter after 24 hours of stimulation with the indicated doses of PMA, showing a binary pattern of activation wherein the frequency of cells responding, but not intensity of the response, increases with the dose.

(D) Histograms of T-Sapphire expression for 293T WT cells containing the NF-κB reporter after 24 hours of stimulation with the indicated doses of TNF-a.

(E) Microscopy images of 293T cells containing the NF-κB reporter after 24 hours of stimulation with the indicated doses of TNF-a. No signal is detected in unstimulated cells. Cells responded with different intensities at an intermediate dose of 500 pg/mL, but responded with uniform intensity at the highest dose.

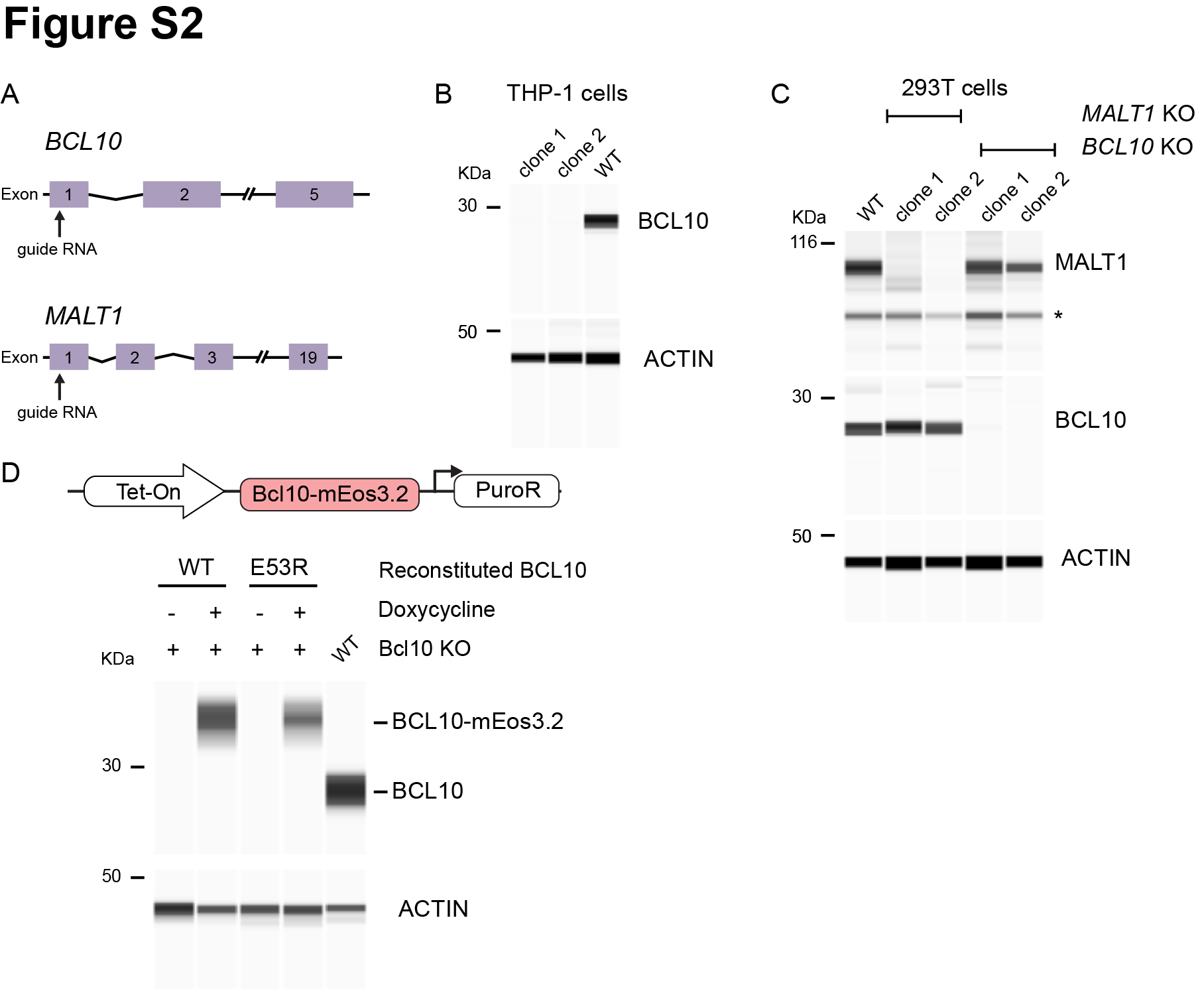

Supplementary Figure 2.

(A) Schema of the *BCL10* and *MALT1* loci indicating the respective sites targeted by guide RNA for CRISPR-Cas9 mediated knockout approaches. After transfection of plasmids containing Cas-9 expression with the respective guide RNA, we verified the correct KO by targeted deep sequencing and western blot.

(B) Capillary immunodetection of BCL10 protein in lysates of THP-1 cells. We selected two independent clones to perform experiments and confirm complete ablation of BCL10 expression after CRISPR-Cas9 mediated KO.

(C) Capillary immunodetection of BCL10 and MALT1 proteins in 293T cells. We selected 2 clones that completely lost the expression of BCL10 and MALT1.

(D) Top, schematic of the lentiviral construct integrated into THP-1 *BCL10*-KO cells to reconstitute the expression of BCL10 fused to mEos3.2. Bottom, capillary immunodetection of BCL10 in cells following doxycycline (1 ug/ml) treatment, showing tight control over BCL10 expression.

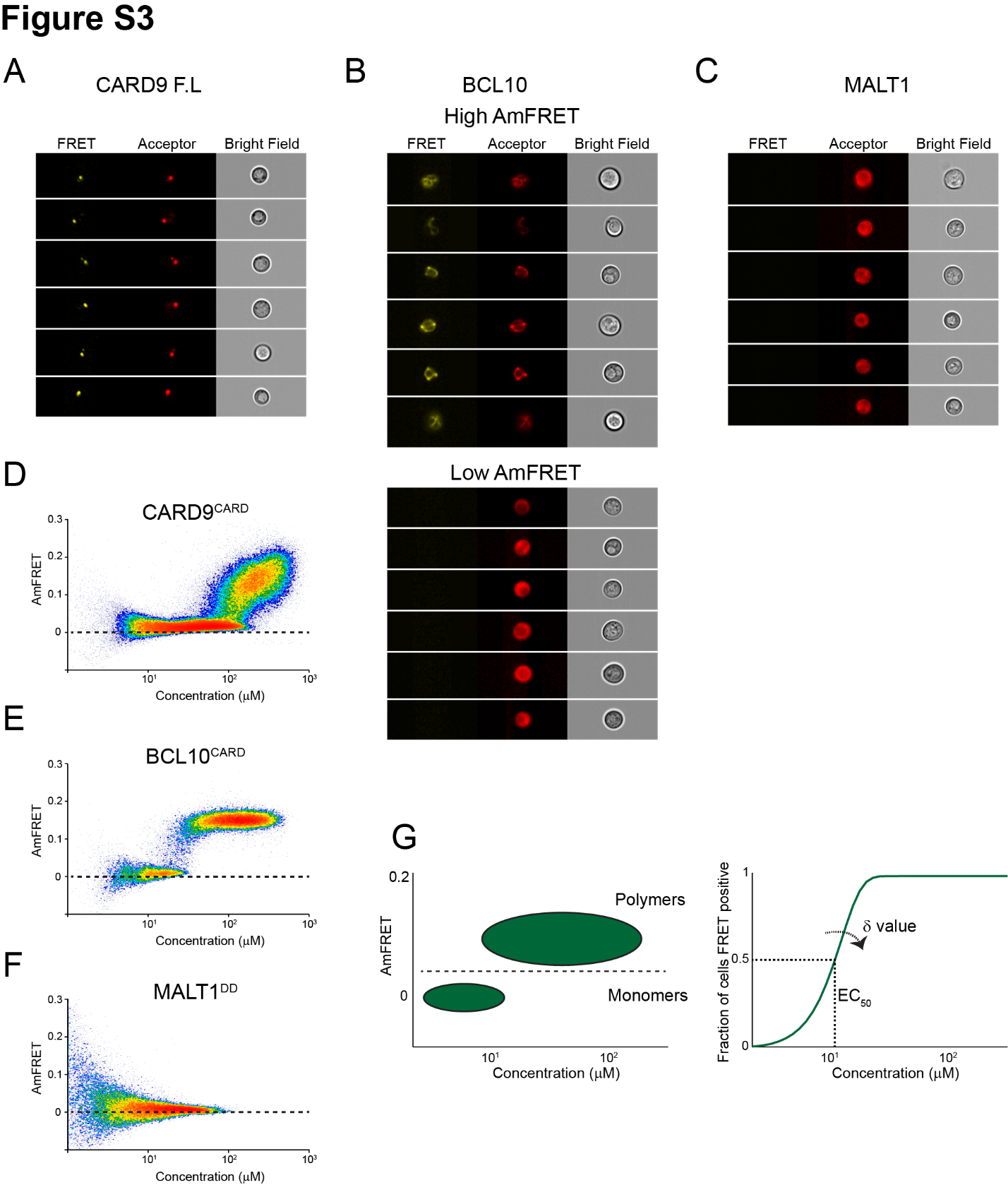

Supplementary Figure 3.

(A) Imaging flow cytometry images from the DAmFRET experiment in Fig 2B, showing that FL CARD9 localizes to a punctum in yeast cells.

(B) Imaging flow cytometry images from the DAmFRET experiment in Fig 2C, showing that FL BCL10 is polymerized in cells with high AmFRET (top), but is diffuse in cells with low AmFRET.

(C) Imaging flow cytometry images from the DAmFRET experiment in Fig 2D, showing a diffuse distribution of FL MALT1.

(D) DAmFRET plot of CARD9^CARD^ showing the onset of polymerization, with only a small discontinuity, at approximately 100 uM.

(E) DAmFRET plot of BCL10^CARD^ showing a discontinuous distribution of cells between low- and high-AmFRET populations, indicating nucleation-limited polymerization.

(F) DAmFRET plot of MALT1^DD^ showing that the protein does not self-associate.

(G) Schematic representation of DAmFRET Weibull fit analysis. DAmFRET plots are gated to define populations of cells containing monomer or polymers. After obtaining the fraction of cells FRET positive across the expression range, we compute the Weibull fit analysis. From this we derive two parameters, EC_50_ and δ. The parameter δ relates to the independence of nucleation on concentration.

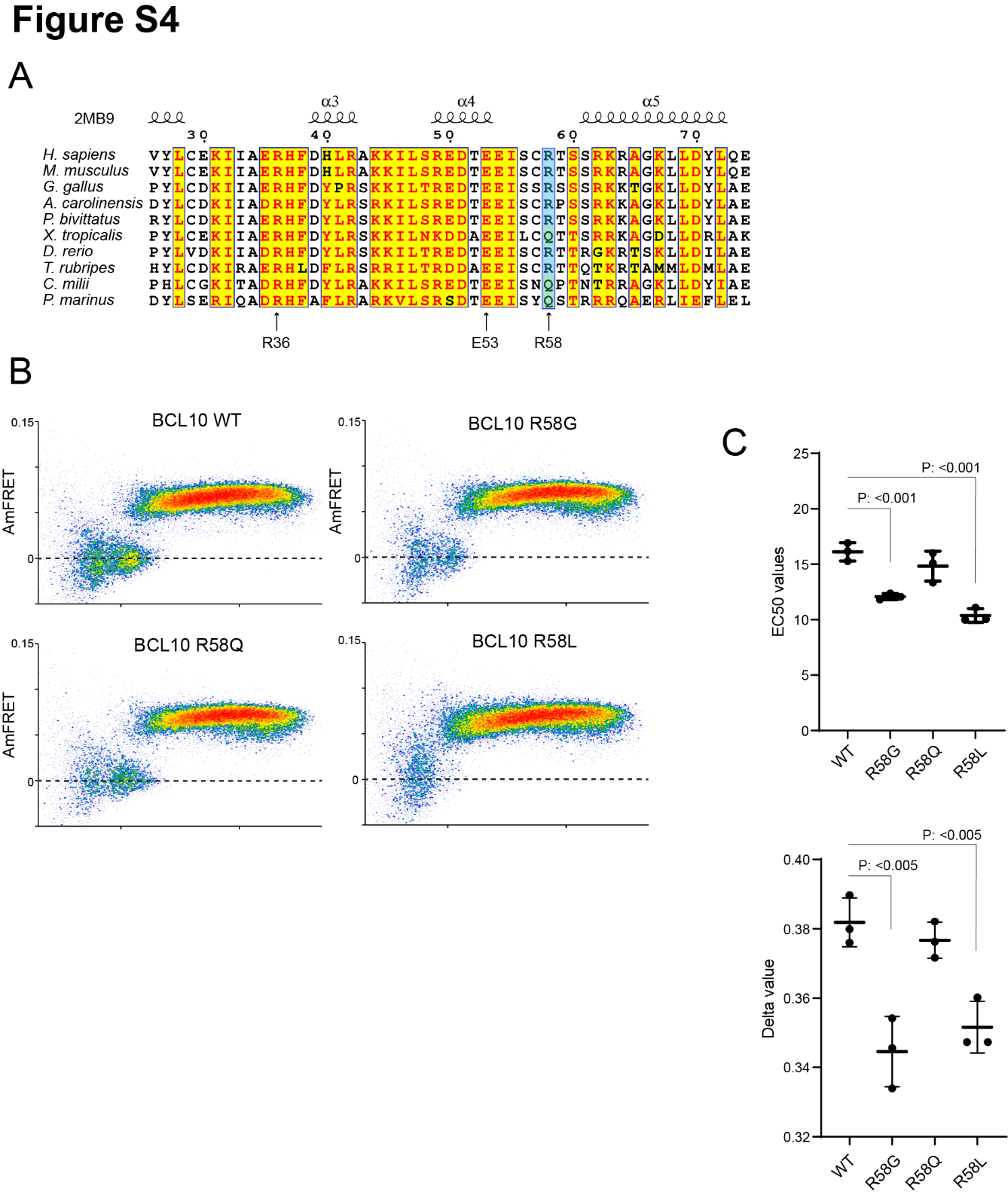

Supplementary Figure 4.

(A) Multiple sequence alignment of vertebrate *BCL10* homologs. Arrows indicate the positions of mutations characterized herein.

(B) DAmFRET plots of yeast cells expressing the indicated mutants of FL BCL10.

(C) EC_50_ (top) and **ẟ** (bottom) values of Weibull fits to DAmFRET plots of BCL10 mutants. Data are from three independent experiments. Comparisons were made with the unpaired t-test.

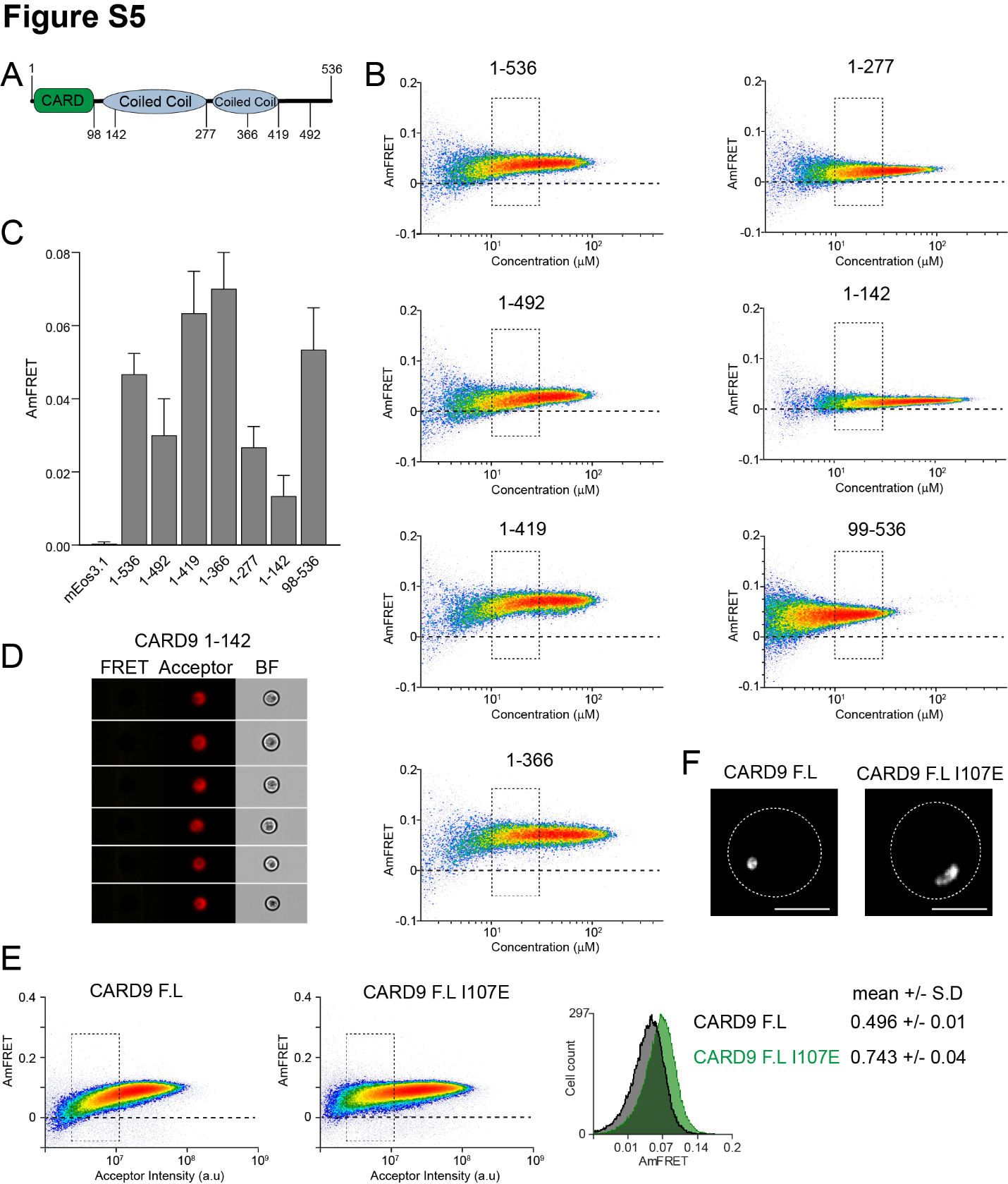

Supplementary Figure 5.

(A) Schematic of the CARD9 protein, indicating the boundaries of truncated versions tested.

(B) DAmFRET plots for truncated versions of CARD9. This analysis reveals the contribution of the coiled-coil domain to the higher-order assembly of CARD9. Even in the absence of the CARD domain, CARD9 still retains comparable FRET levels as the full-length version.

(C) Median AmFRET values from 4 independent replicates of the region delimited in plots in (B).

(D) Imaging flow cytometry images show that CARD9^1-142^ is diffuse despite appreciable AmFRET, consistent with its expected dimerization.

(E) DAmFRET plots of CARD9 FL and CARD9 I107E show that the I107E mutation increases assembly at low concentrations. Inset shows the histogram of AmFRET values obtained from the gates shown in the DAmFRET plots.

(F) Representative images of cells from confocal microscopy showing that the I107E mutant forms relatively irregular puncta.

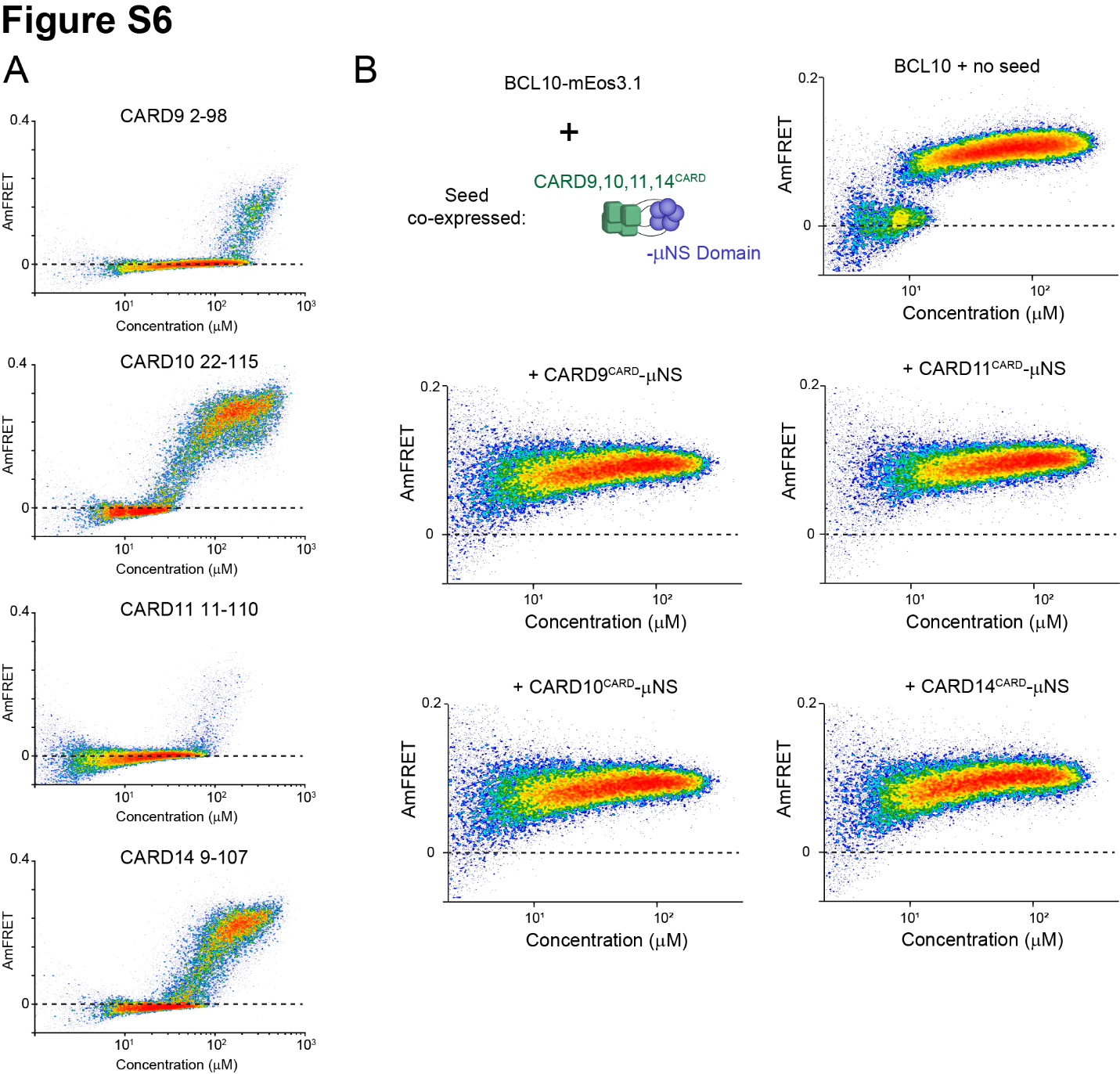

Supplementary Figure 6.

(A) DAmFRET plots for the CARD domains of the CARD-CC family members, CARD9, CARD10, CARD11, and CARD14, showing that all four polymerize in a concentration-dependent fashion with a negligible nucleation barrier.

(B) Schematic and DAmFRET plots showing the nucleation of BCL10-mEos3.1 by the CARD domains of either CARD9,10,11 or 14 fused to μNS. All CARD domains eliminated the low-AmFRET population of cells expressing BCL10-mEos3.1.

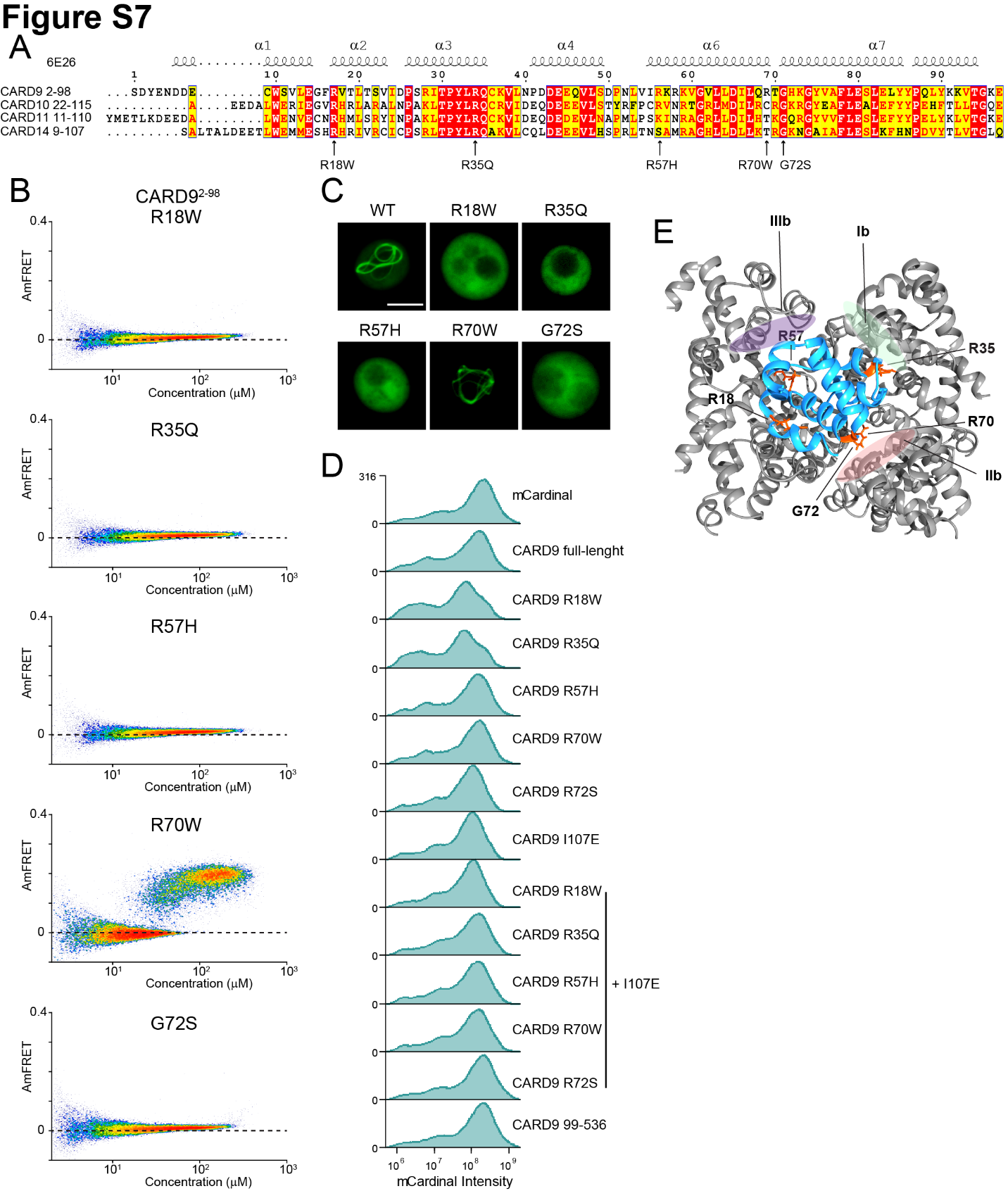

Supplementary Figure 7.

(A) Sequence alignment of the human CARD-CC family members. Arrows indicate pathogenic mutations in CARD9.

(B) DAmFRET plots for CARD9^CARD^ mutants, related to the analysis in figure 2H.

(C) Fluorescence microscopy images of yeast cells expressing CARD9^CARD^-mEos3.1 showing that the pathogenic mutations, with the exception of R70W, disrupt assembly. Scale bar 5 µm.

(D) Histograms of mCardinal fluorescence intensity corresponding to the expression levels of CARD9 mutants 48 hours after transfection into 293T NF-κB reporter cells.

(E) Structure of the CARD9^CARD^ polymer (6N2P) highlighting the residues (orange) whose mutation causes susceptibility to fungal infections, with respect to polymer interfaces.

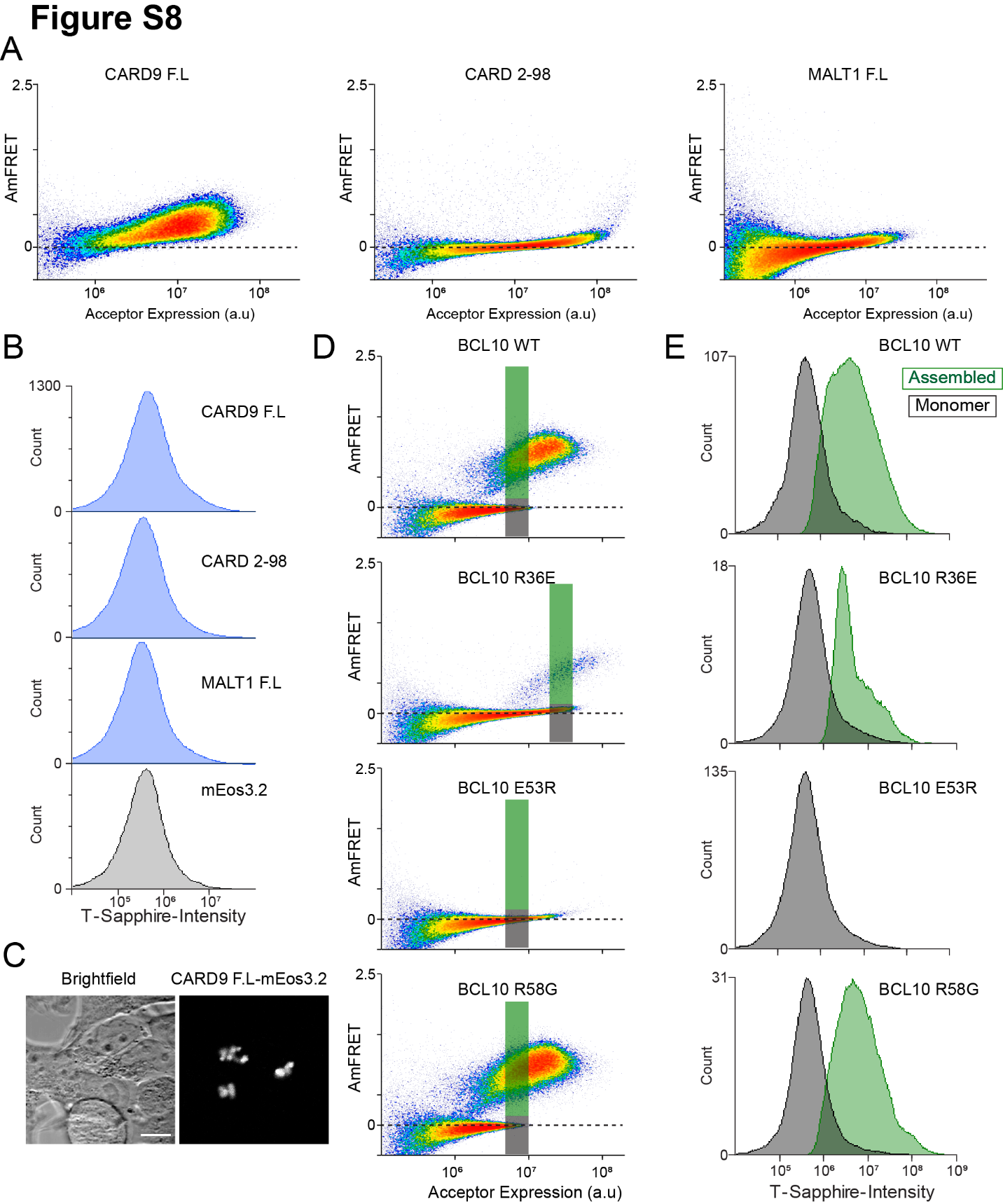

Supplementary Figure 8.

(A) DAmFRET plots of CARD9 full-length, CARD9^CARD^, or MALT1 full-length tagged with mEos3.2 48hrs post-transfection in 293T NF-κB reporter cells.

(B) Histograms of T-Sapphire fluorescence in the 293T NF-κB reporter cells expressing the indicated proteins. Neither CARD9 nor MALT1 were able to induce activation of NF-κB.

(C) Microscopy images of CARD9-mEos3.2 expressed in 293T cells, showing puncta formation even in the absence of stimulation.

(D) DAmFRET plots of 293T *BCL10*-KO cells containing the NF-κB reporter 48 hrs after transfection with the indicated *BCL10* constructs. The green and gray rectangles indicate the gated regions evaluated for T-Sapphire intensity in (E).

(E) Histograms of T-Sapphire fluorescence for cells expressing the same concentration of BCL10 in either a monomer (gray) or polymerized (green) form.

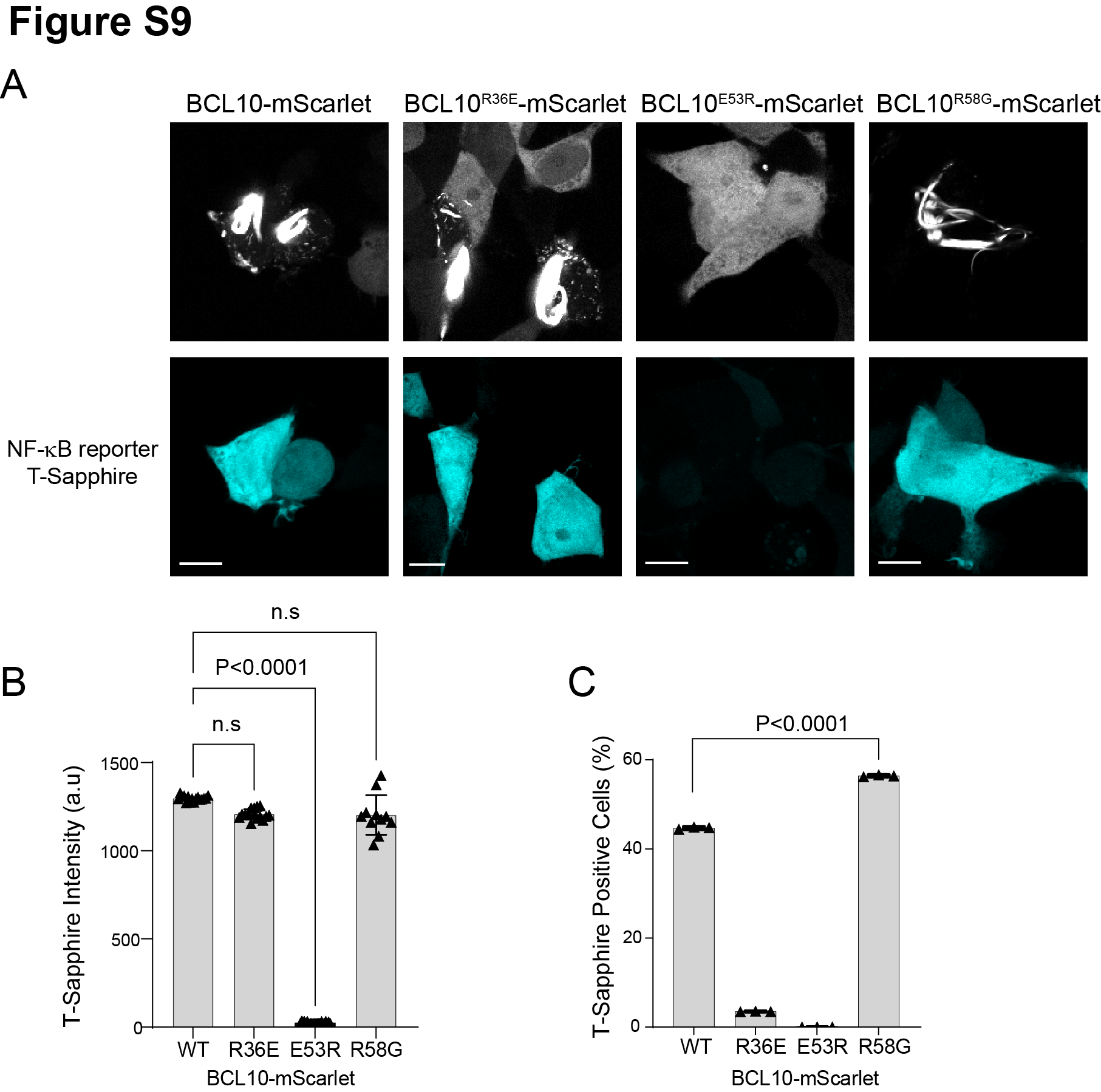

Supplementary Figure 9.

(A) Confocal microscopy images of 293T *BCL10*-KO cells containing the NF-κB 48hrs after transfection with the indicated *BCL10* constructs, showing that cells containing BCL10 polymers express T-Sapphire, indicating NF-κB activation. Scale bar 10 µm.

(B) T-Sapphire intensity normalized by cell area for each of the corresponding *BCL10* constructs. This analysis includes >30 cells per sample from 3 independent experiments. T-Sapphire is expressed to the same level in all cells with polymers, reflecting the binary activation of NF-κB.

(C) Percentage of cells with T-Sapphire expression. For this analysis, we included 4 randomly selected imaging fields from 3 independent experiments. The number of cells quantified for each group contains >300 cells. Two sample comparisons were performed using unpaired t-test.

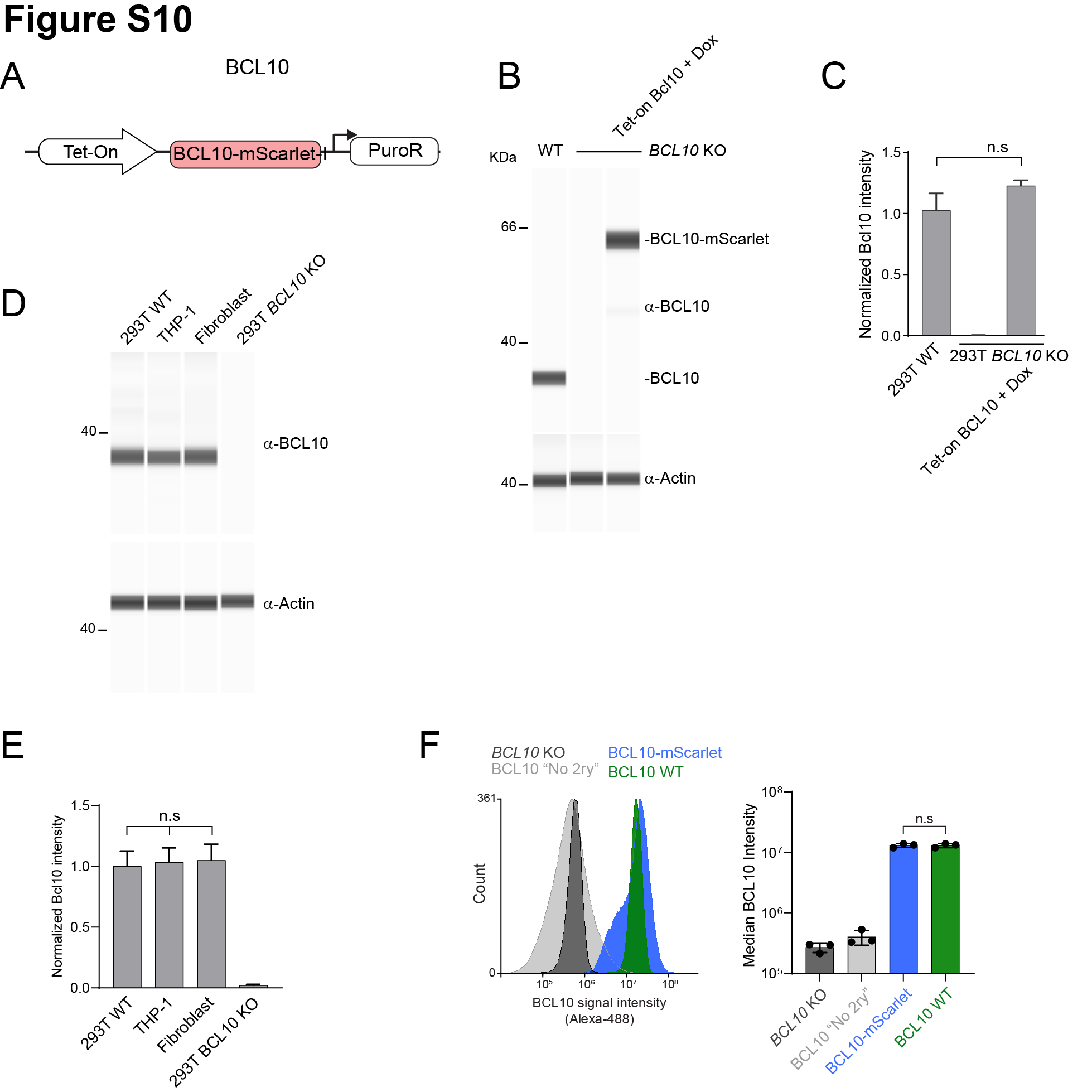

Supplementary Figure 10.

(A) Schematic of the lentivirus construct to reconstitute BCL10 expressed as a fusion to -mScarlet.

(B) Capillary immunodetection of BCL10 levels in WT, KO, and reconstituted 293T cells. The latter were stimulated with 1 ug/ml Dox for 24 hours. Lysates were generated from 2 million cells in each case.

(C) Quantification of BCL10 protein levels from three independent experiments as shown in (B). BCL10 intensities were normalized to that of actin in each lane. Reconstituted cells express approximately endogenous levels of BCL10, fused to mScarlet. Statistical comparisons were made using unpaired t-test.

(D) Capillary immunodetection of BCL10 in 293T WT cells, THP-1 cells, and primary human fibroblasts. Lysates were generated from 1 million cells in each case.

(E) Quantification of BCL10 protein levels from three independent experiments as shown in (D). BCL10 is expressed to the same level in these diverse human cells. Statistical comparisons were made using unpaired t-test.

(F) Left, flow cytometry histograms of anti-BCL10 staining for the indicated cell lines. 293T *BCL10*-KO cells reconstituted with *BCL10*-mScarlet were stimulated with Dox 1 ug/ml for 24 hours. Right, median BCL10 intensity of three independent experiments. Wildtype and BCL10-mScarlet expression levels are highly similar.

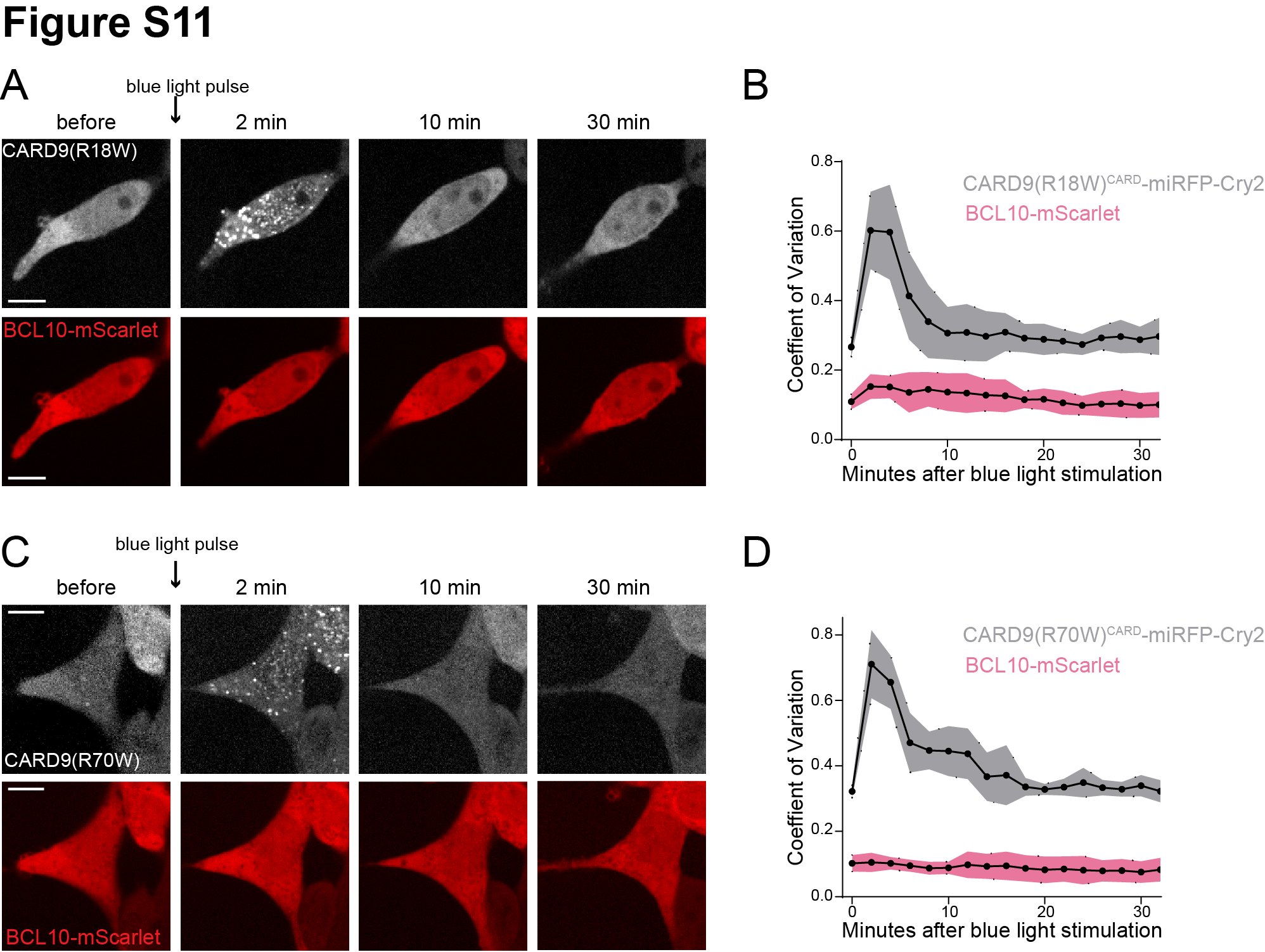

Supplementary Figure 11.

(A) Time-lapse microscopy of 293T *BCL10-*KO cells stably expressing CARD9(R18W)^CARD^-miRFP670nano-Cry2clust and BCL10-mScarlet. Cells were maintained at 37°C with 5% CO_2_. Images were taken every 2 minutes following 1 second exposure of 488 nm light stimulation. Scale bar 10 µm.

(B) Values of the coefficient of variation of mScarlet and miRFP670nano pixel intensities over time. The data are from 20 cells from independent experiments.

(C) Time-lapse microscopy of 293T *BCL10*-KO cells stably expressing CARD9(R70W)^CARD^-miRFP670nano-Cry2clust and BCL10-mScarlet. Images were taken every 2 minutes following a short pulse of 488 nm light stimulation. Scale bar 10 µm.

(D) Values of the coefficient of variation of mScarlet and miRFP670nano pixel intensities over time. The data are from 20 cells from independent experiments.

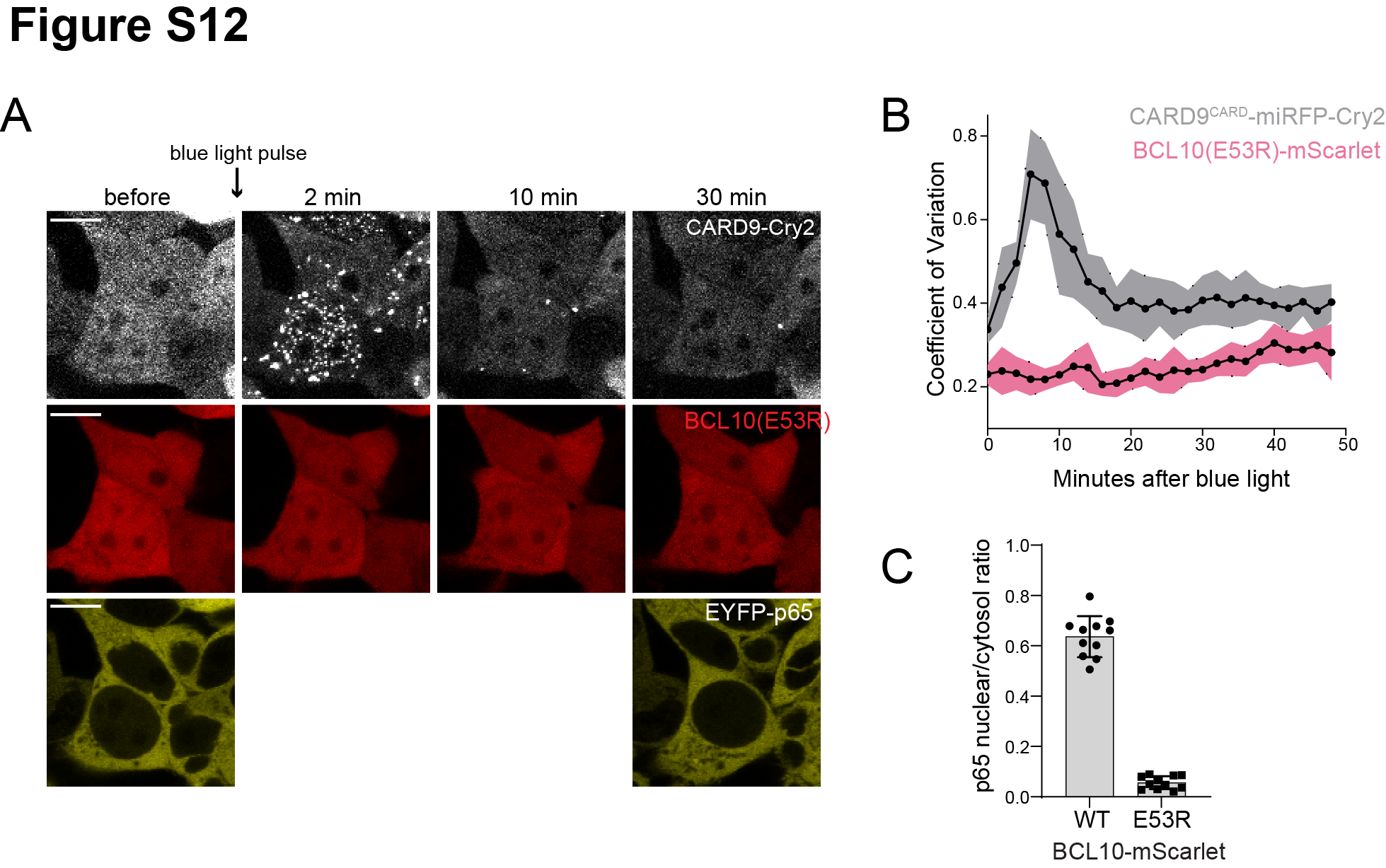

Supplementary Figure 12.

(A) Time-lapse microscopy of 293T *BCL10*-KO cells stably expressing CARD9^CARD^-miRFP670nano-Cry2clust, BCL10(E53R)-mScarlet, and EYFP-p65. Cells were maintained at 37°C with 5% CO_2_ and imaged every 2 minutes following a short pulse of 488nm light stimulation. Scale bar 10 µm. To avoid undesired Cry2clust activation, EYFP-65 was imaged only at the beginning and end of the time course.

(B) Values of the coefficient of variation of mScarlet and miRFP670nano pixel intensities over time. The data are from 20 cells from independent experiments.

(C) Ratios of EYFP fluorescence intensities in the nucleus versus cytosol 30 minutes after blue light stimulation in (A). The quantification included at least 10 cells per group.

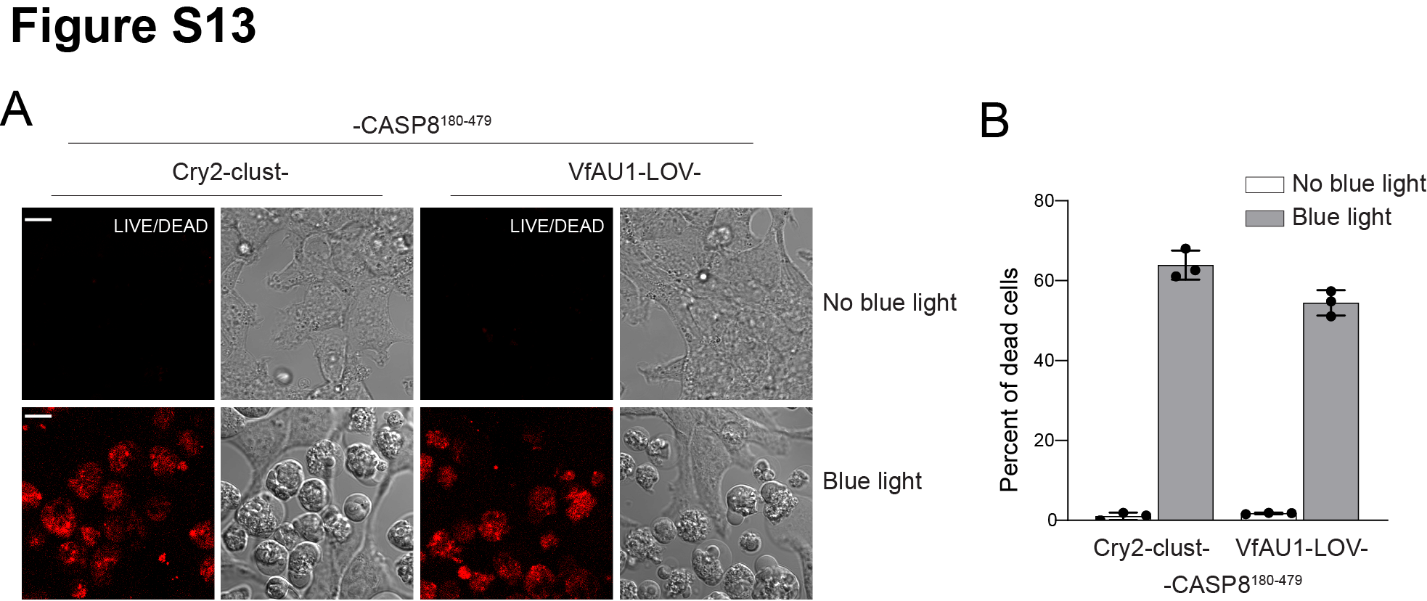

Supplementary Figure 13.

(A) Microscopy images of 293T cells transfected with either Cry2clust or VfAU1-LOV fused to CASP8^180-479^, which lacks the death domain. One day after transfection, cells were exposed to constant blue light stimulation from a blue LED source while being maintained at 37°C and 5% CO_2_. Prior to imaging, cells were treated with the LIVE/DEAD cell marker DRAQ7.

(B) Percentage of dead cells from the experiment in A. Data are from 3 independent experiments with >200 cells counted per replicate. Scale bar 10 µm.

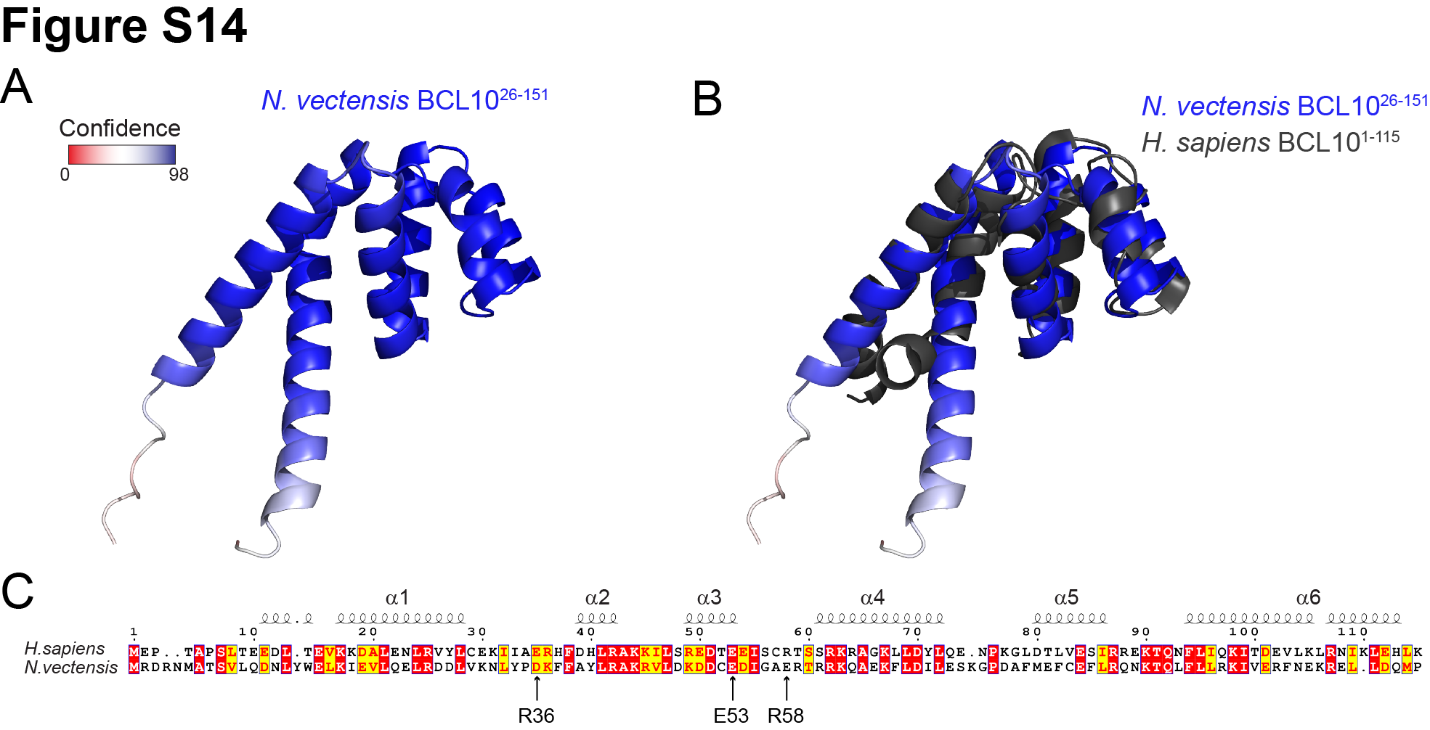

Supplementary Figure 14.

(A) AlphaFold structure of *Nematostella vectensis* BCL10 (A7SSM3). The selected model shows a very high degree of confidence throughout the CARD domain.

(B) Alignment of the predicted structure of Nv BCL10 with human BCL10 (2MB9). The structural alignment produces an RMSD of 2.912.

(C) Sequence alignment for the CARD domain of Nv and human BCL10. Arrows indicate the mutations in human BCL10 that perturb nucleation.
